# Multimodal single-cell profiling reveals cancer crosstalk between macrophages and stromal cells in poor prognostic cholangiocarcinoma patients

**DOI:** 10.1101/2024.02.03.578669

**Authors:** Lara Heij, Sikander Hayat, Konrad Reichel, Sidrah Maryam, Colm J. O’Rourke, Xiuxiang Tan, Marlous van den Braber, Jan Verhoeff, Maurice Halder, Fabian Peisker, Georg Wiltberger, Jan Bednarsch, Daniel Heise, Julia Campello Deierl, Sven A. Lang, Florian Ulmer, Tom Luedde, Edgar Dahl, Danny Jonigk, Jochen Nolting, Shivan Sivakumar, Jens Siveke, Flavio G. Rocha, Hideo A. Baba, Jesper B. Andersen, Juan J. Garcia Vallejo, Rafael Kramann, Ulf Neumann

**Affiliations:** Department of Surgery and Transplantation, University Hospital RWTH Aachen, Aachen, Germany; Department of Surgery and Transplantation, University Hospital Essen, Essen, Germany; Department of Pathology, University Hospital Essen, Essen, Germany; Department of Renal and Hypertensive Disorders, Rheumatological and Immunological Diseases (Medical Clinic II), Medical Faculty, RWTH Aachen University, Aachen, Germany; Amsterdam UMC – Location Vrije Universiteit Amsterdam, Molecular Cell Biology & Immunology, De Boelelaan 1117, Amsterdam, The Netherlands; Biotech Research and Innovation Centre (BRIC), Department of Health and Medical Sciences, University of Copenhagen, Denmark; Amsterdam Infection & Immunity, Cancer Immunology, Amsterdam, The Netherlands; Tytgat Institute for Liver and Intestinal Research, Amsterdam UMC, University of Amsterdam, Amsterdam, the Netherlands; Department of Surgery, Maastricht University Medical Center (MUMC), Maastricht, The Netherlands; Department of Gastroenterology, Hepatology and Infectious Diseases, University Hospital Duesseldorf, Düsseldorf, Germany; Institute of Pathology, University Hospital RWTH Aachen, Aachen, Germany; Center for Integrated Oncology Aachen Bonn Cologne Duesseldorf (CIO ABCD), D-52074 Aachen, Germany; Biomedical Research in Endstage and Obstructive Lung Disease Hannover (BREATH), German Center for Lung Research (DZL), 30625 Hannover, Germany; Institute of Pathology, Hannover Medical School, 30625 Hannover, Germany; University of Birmingham and University hospitals of Birmingham NHS Trust, Birmingham, B15 2TT, UK; Bridge Institute of Experimental Tumor Therapy, West German Cancer Center, University Hospital Essen, University Duisburg-Essen, Essen, Germany; Division of Surgical Oncology, Knight Cancer Institute, Oregon Health and Science University, Portland, OR, USA

**Author notes:** Denotes joint first authors (Lara Heij, Sikander Hayat, Konrad Reichel). denotes joint last authors (Juan J. Garcia Vallejo, Rafael Kramann, Ulf Neumann). Corresponding author: Lara Heij Department of Surgery and Institute of Pathology University Hospital Essen Hufelandstrasse 15, Essen. Author contributions: LH, SH, KR, JJGV, RK, and UN designed this study. SL, FU, XT, GW, JB, MB, FP, DH, JN, JV, and MH contributed to sample collection and processing. LH, SH, KR, SM, CR, and JJGV were responsible for data analysis and interpretation. LH, DJ, and HB provided histological expertise. LH, SH, SM, RK, JA, JD, and CR performed statistical analysis and interpretation. LH, SH, SS, FR, JJGV, JA, CR, and KR contributed to the first draft of this manuscript. UN, RK, SS, TL, DJ, ED, JS, HB, and JJGV provided the infrastructure and supervised the study. All authors contributed to the data analysis and manuscript writing. Data availability: The code used in analyzing this data is available at https://github.com/hayatlab/cholangiocarcinoma_ici. The data will be made available via https://zenodo.org/uploads/10017884.

**Keywords:** Cholangiocarcinoma, Tumor Microenvironment, Transcriptomics, Immune Exhaustion

## Abstract

**Background and aims:** Cholangiocarcinoma (CCA) is a deadly disease, and this cancer entity is characterized by an abundant stroma. The tumor microenvironment (TME) plays an important role in aggressive behavior and poor response to therapeutics; however, underlying pathways are unknown.

**Methods:** To fill this gap, we used multiplexed immunohistochemistry, high-dimensional phenotyping, and transcriptomics to analyze human CCA samples and identify cell cluster crosstalk in patients with a poor prognosis.

**Results:** Our findings confirmed the presence of Tregs and the lack of effector memory cells in the tumor. New findings are the spatiality of the effector memory cells being more present in the peripheral tissue, for some reason these immune cells fail to reach the tumor niche. We revealed cancer crosstalk with macrophages and stromal cells and identified responsible genes in the poor prognosis group. Amongst the responsible ligand pairs are GAS6-AXL belonging to the TAM family. We then identified VCAN-TLR2 to be present and influencing the ECM in a way to supports immune exhaustion. Last, EGFR-TGF-β is expressed in macrophages and this finding is important in Tregs induction and blocking cytotoxic T cell function.

**Conclusions:** The multiple mechanisms leading to the exclusion of relevant immune cells needed for an anti-cancer response and mechanisms leading to active immune suppression are part of complex cell-cell crosstalk. This study provides a deeper insight into the immune exhausted phenotype in CCA.

## Introduction

Cholangiocarcinoma (CCA) is a deadly disease occurring along the biliary tract and is considered a rare cancer type with a 5-year overall survival (OS) rate of 7–20%(1, 2). CCA is classified as intrahepatic cholangiocarcinoma (iCCA), perihilar cholangiocarcinoma (pCCA), or distal cholangiocarcinoma (dCCA). Although the histological pattern can be very similar, the subtypes have been considered different entities because of differences in their surrounding tissue and molecular tumor heterogeneity(3, 4, 5). CCA typically shows distinctive perineural invasion (PNI), defined as the invasion of cancer cells into and around the nerve trunks. Recently, we showed that both iCCA and pCCA may present a high nerve fiber density (NFD) in the tumor microenvironment (TME), which is linked to better survival in a subset of patients(6, 7). Another shared histological feature of CCA subtypes is the presence of an abundant stromal compartment. The stroma displays heterogeneous cell types, including immune cells and fibroblasts. Interactions between different cell types result in immune-infiltrated or immune-excluded phenotypes(8, 9, 10).

Recent progress in cancer care has included the development of immune checkpoint inhibitors (ICI) and therapeutic agents that lead to host immune cell recruitment to target cancer. Specifically, T cells play an important role in the immune response of the host to cancer and are the main factor leading to curative responses to checkpoint inhibitor immunotherapy. In CCA, regulatory T cells (Tregs) dominate the early immune response and remain present in later tumor stages(11, 12), whereas effector T cells are rarely found, and if present, do not show signs of activation(13). The presence of CD4 and CD8 T cells infiltrating the tumor and peritumoral areas is a sign of better outcomes(14), and high numbers of CD8 T cells in the outer border of the tumor are positively correlated with 5-year survival rates in patients with iCCA(15). Furthermore, a subset of CD8 T cells expressing CD161 is known to have higher cytotoxic potential(16, 17), and high levels of CD161 expression are associated with protective functions in many cancer types, including CCA(18). The immune checkpoint PD-1 is known to promote immune evasion, and high PD-1 levels in tumor-infiltrating CD8 T cells in iCCA were previously found to be associated with an unfavorable prognosis(19). Nevertheless, our understanding of the immune cell environment in CCA remains limited, and additional factors related to treatment responses are still needed(20).

In addition to T cells, tumor-associated macrophages (TAMs) constitute another major immune population within the TME. Although TAMs can exhibit both pro- and anti-inflammatory functions, they are thought to be mostly immunosuppressive(21). Despite some disagreement, most studies point to a worse prognosis in patients with high levels of TAMs (22, 23, 24). TAMs are a major source of PD-1 ligand (PD-L1) in CCA tumors(25). Furthermore, TAMs can secrete cytokines, such as tumor growth factor-β (TGF-β), which suppresses the T cell response by T cell exclusion and prevents the acquisition of the tumor-suppressive TH1-effector phenotype(26). Despite its immune-regulatory functions, modulation of the extracellular matrix (ECM) plays a crucial role in recruiting TAMs in tumors(27).

In this study, we aimed to better characterize the stromal compartment in CCA through a comprehensive approach using cytometry by time-of-flight (CyTOF) and single nuclei (sn)RNAseq. We focused on patients with primary resected iCCA and pCCA and characterized the differences in immune landscapes of both entities in primary tumors, compared to matched peripheral liver tissue, and peripheral blood samples. To identify the crosstalk between cell clusters responsible for the immunosuppressive environment, we used snRNA sequencing to characterize the cellular heterogeneity of the transcriptome in patients with pCCA. We identified signs of exhaustion and an increased abundance of regulatory T cells in the tumor niche, while CD8 effector memory (EM) and tissue-resident effector memory cells (TRMs) failed to reach the tumor and were restricted to the peripheral liver tissue. To further explore the immunosuppressive TME, we revealed a shift in crosstalk between stromal cells and specific cancer cells, mainly present in the poor prognosis group. In addition, these cancer cells demonstrated an upregulated cell communication with macrophage-myeloid cells. Overall, this study sheds light on the etiology of immunosuppressive TME in CCA.

## Results

To elucidate the heterogeneity of the TME, we performed high-dimensional immunophenotyping using mass cytometry time-of-flight (CyTOF) on 16 tumor samples. Of these, seven were iCCA patients and nine were pCCA patients with resectable disease. For all patients, we obtained matched tumor-free liver tissue (hereafter referred to as ‘peripheral tissue’), central tumor tissue, and peripheral blood samples (Figure 1). All patients underwent surgery at the University Hospital RWTH Aachen to obtain detailed patient characteristics (see Supplementary Table 1). Mass cytometry was performed using the panel described in Supplementary Table 2, with a strong emphasis on the characterization of different subsets of T cells and the expression of known costimulatory molecules and immune regulators of importance in cancer immunotherapy. This was later combined with immunohistochemistry (IHC) to give more spatial context and results from snRNA-seq to gain a more complete understanding of the TME (Figure 1).

**Figure 1.**
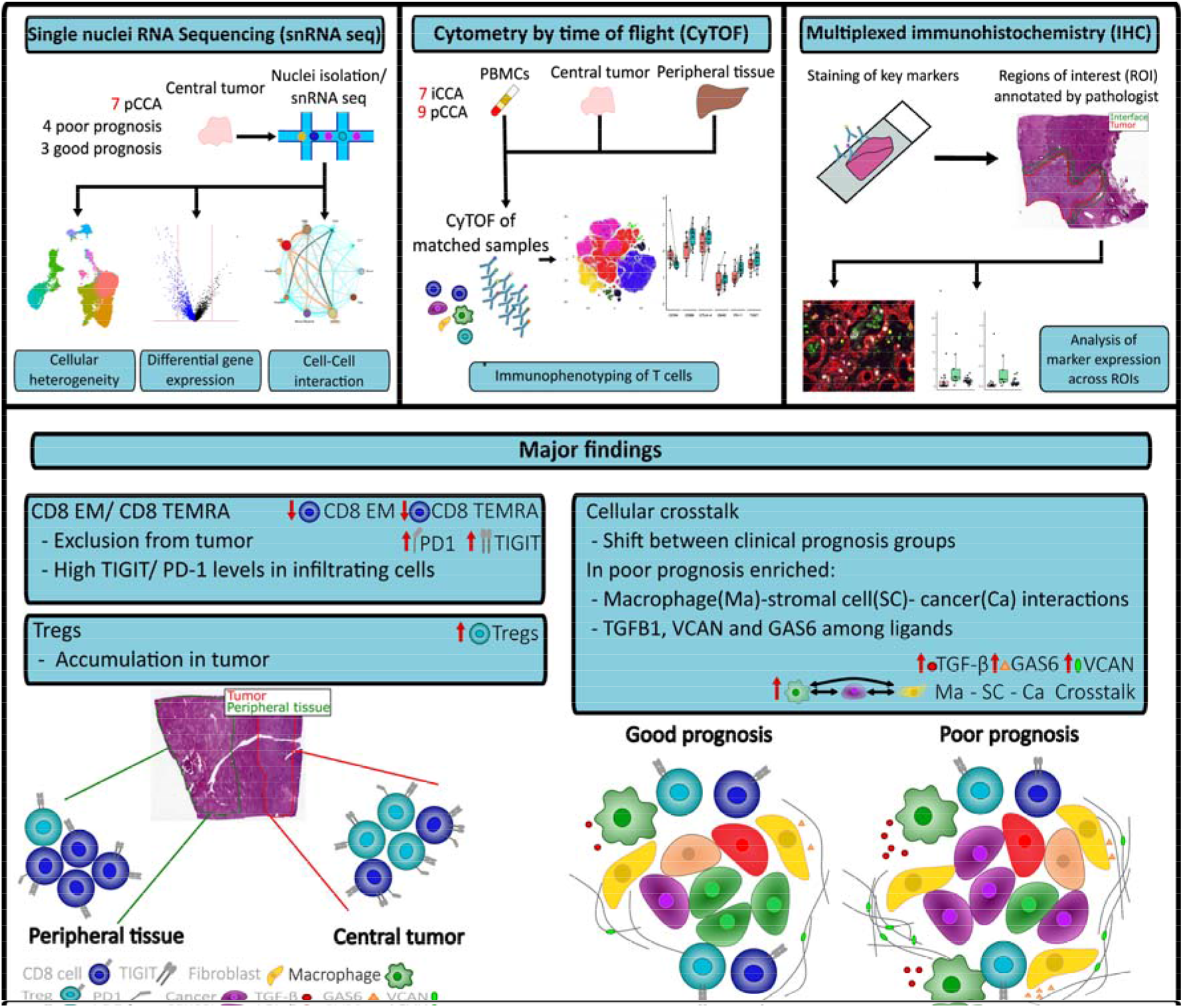
Study workflow and overview of results. In our study, we included patients with primary intrahepatic cholangiocarcinoma (iCCA) or perihilar cholangiocarcinoma (pCCA) scheduled for resection. To examine the tumor microenvironment (TME) we used single nuclei RNA sequencing (snRNAseq), Cytometry by time of flight (CyTOF) and multiplexed immunohistochemistry (IHC). For snRNAseq, we analyzed 7 pCCA samples grouped into poor prognosis (n = 4) and good prognosis (n = 3) groups based on clinical data. For CyTOF we used matched blood, tumor, and peripheral samples from 7 iCCA and 9 pCCA patients. To gain further spatial information we used IHC data where a pathologist annotated tumor/ interface regions. Combining the data, we were able to identify an exclusion of CD8 T cell subsets from the tumor and an accumulation of regulatory T cells (Tregs) in the tumor. Tumor-infiltrating T cells expressed high levels of several immune checkpoints. We also revealed a shift in the cell-cell interactions in the poor prognosis patients. This shift was marked by increased interactions between macrophages, stromal cells, and cancer cells. Enriched ligands included VCAN, TGFB1, and GAS6.

First, we used a combination of unsupervised clustering and manual gating to identify T cells in the CyTOF data and focused on the analysis of CD4 and CD8 cells (Supplementary Figure S1). Downstream analysis of both compartments using a combination of dimensionality reduction (UMAP) and clustering (Phenograph) resulted in the identification of 27 CD4 T-cell clusters and 31 CD8 T-cell clusters, which were manually curated and annotated into known immune cell subsets (Supplementary Figure S2).

### The spatial distribution of CD4 and CD8 memory cells differs between peripheral tissue and tumor tissue, but not between tumor entities

Using an unsupervised clustering-based approach, we uncovered differences between the CD4 and CD8 memory compartments. Notably, the abundance of none of the examined subsets was significantly elevated when comparing the iCCA and pCCA tumor samples (Figure 2a). This suggests a comparable T-cell composition in the iCCA and pCCA subtypes, which enabled a combined analysis of both entities. Looking at cell abundance between the matched tumor and peripheral tissue, we observed a 5-fold increase in CD4 Tregs (p=0.002) and an increase in CD4 tissue-resident memory cells in the tumor tissue (p=0.0003). In the CD8 compartment, CD8 effector memory and tissue-resident effector memory RA cells (TEMRA) were more abundant (p=0.04 and p=0.02, respectively) in the peripheral tissue, and although CD8 tissue-resident memory cell levels were elevated in the tumor tissue, the difference was not significant (Figure 2b). Besides the differences in subset abundance, the low-dimensional representation also revealed a more complex heterogeneity within most subsets (Figure 2c).

**Figure 2.**
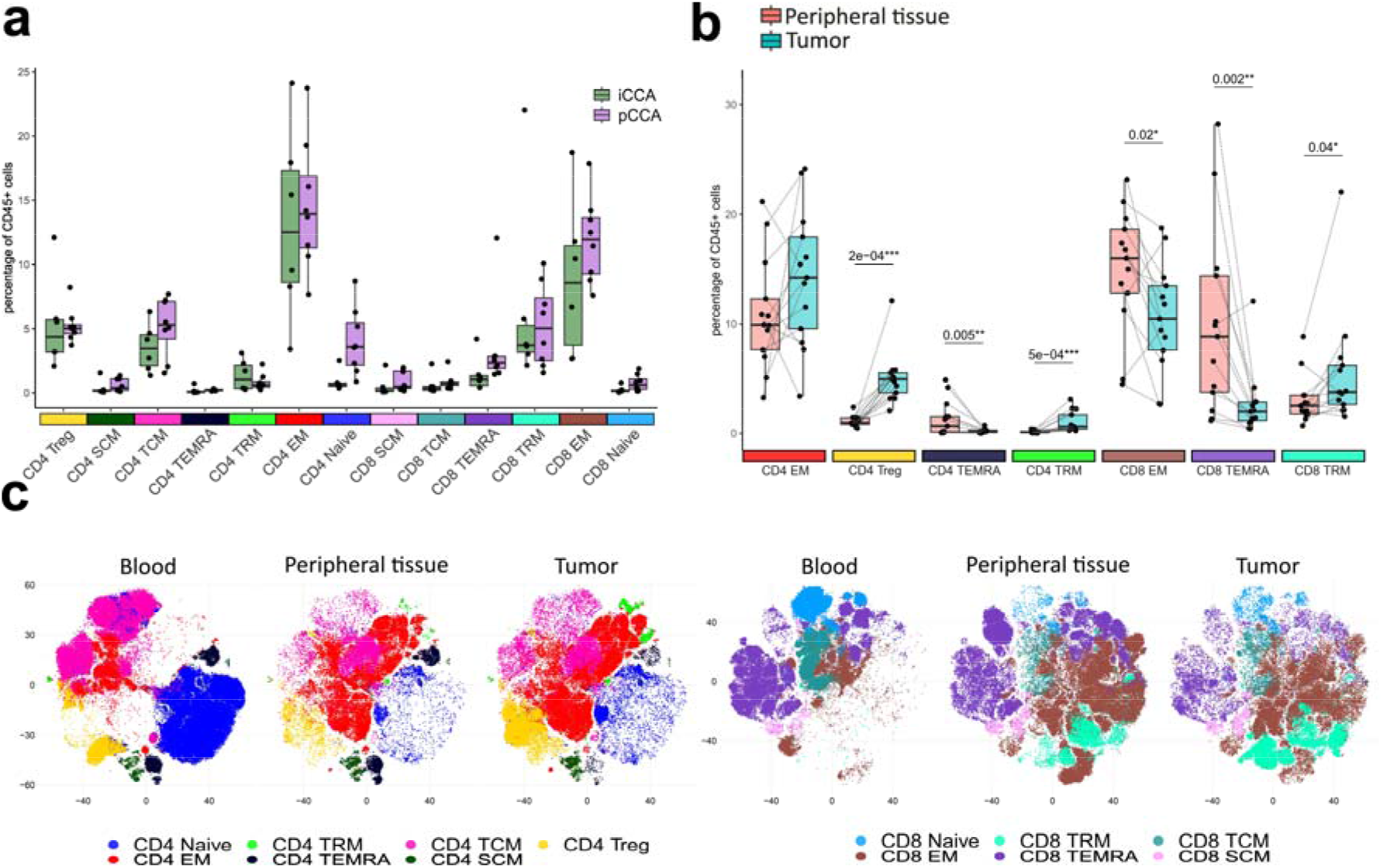
High-dimensional phenotyping in iCCA. **(a)** Mean percentage of T cell subsets of total viable CD45+ cell types in the tumor. Annotation was done by manual annotation of clusters after unsupervised clustering. No subset is significantly enriched in one of the two CCA subtypes. **(b)** The mean percentage of T cell subsets of total viable CD45+ cell types in matched peripheral tissue and tumor samples shows enrichment of Tregs and exclusion of CD8 EM and CD8 TEMRAs from the tumor. Shown p values were obtained using the Wilcoxon test and were adjusted for multiple testing. **(c)** Dimensionality reduction plots (optSNE) of CD4 and CD8 T cells from all patients reveal heterogeneity within different sample locations. Split by origin tissue for CD4 cells (left) and CD8 cells (right).

### Exhausted effector cells in the tumor niche with less cytotoxic potential

In addition to the sparseness of CD8 effector memory (EM) and TEMRA cells in the tumor, CD8 EM showed a decrease in CD161 expression compared to the peripheral tissue, indicating less cytotoxic activity (Figure 3a). In addition, many of the identified memory subtypes display higher levels of immune checkpoints in tumors. While PD-1 and ICOS levels were elevated in CD8 EMs, CD8 TRMs, CD4 TRMs, and CD8 TRMs, the CD4 TRMs also exhibited higher levels of TIGIT. In addition to the elevated levels of immune checkpoints, we also observed an increase in CD57 expression. CD57 is used as a marker for differentiation/senescence and may promote resistance to ICI in cancer (28, 29). Furthermore, the marker expression profile in the small subset of CD4 stem cell memory (SCM) also showed signs of overactivation in tumors with elevated levels of HLA-DR (p<0.01), CTLA-4 (p<0.01), and ICOS (p<0.05).

**Figure 3.**
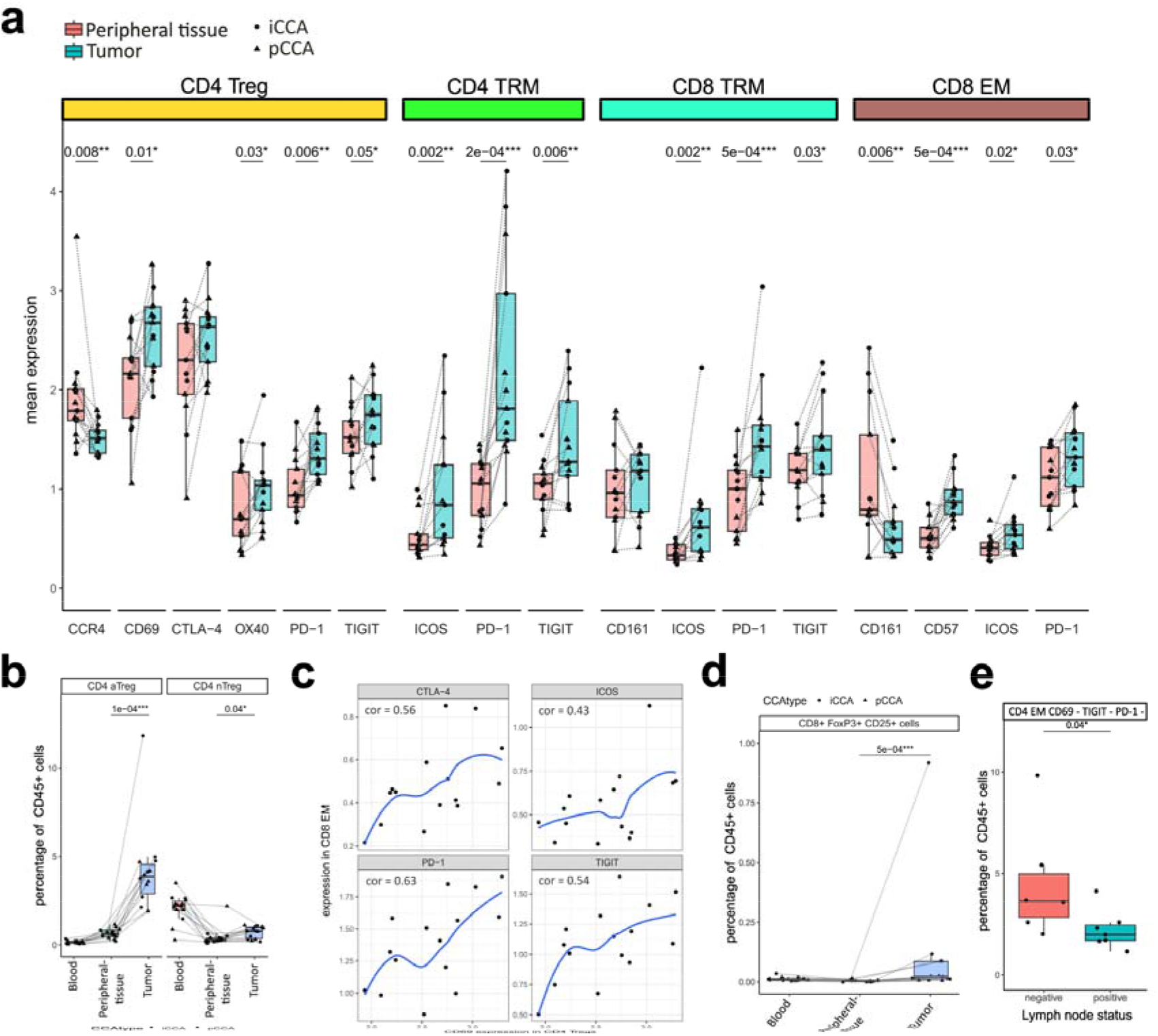
Characterization of marker expression in cell clusters. **(a)** Mean normalized expression of markers across CD4 Treg, CD8 TRM, and CD8 EM cells shows an increased expression of checkpoints/ activations markers in the tumor. **(b)** A specific cluster of T cells (aTreg = activate Treg) with high levels of activation markers is enriched in the tumor but is absent from the blood. Tregs lacking activation markers are also enriched in the tumor but are also common in the blood (nTreg = non-activated Treg). **(c)** Mean checkpoint marker expression in CD8 EM cells correlates with mean CD69 expression in the Treg population within the tumor. The correlation coefficient was calculated using spearman and shown in blue is a regression line. **(d)** CD25 and FoxP3 positive CD8 cells were identified in the tumor by sequential gating. Shown is the mean percentage of total CD45 positive live cells per sample. **(e)** One cluster of CD4 EM cells lacking expression of activation markers and checkpoints is enriched in the peripheral tissue of CCA patients without cancer cell invasion of adjacent lymph nodes.

### Activated Regulatory T cells correlate with exhausted CD8 effector memory cells

CD4 Tregs showed elevated levels of the activation markers CD69 and OX40, and the immune checkpoints PD-1, CTLA-4, and TIGIT. The chemokine receptor CXCR3 (p value did not reach significance) and CCR4 showed decreased levels in the tumor. Both receptors are associated with the migration to tumor tissue(30). Within the total Treg population, we identified a distinct cluster of Tregs with lower levels of CCR7 and higher levels of CD69, indicating a more active state (Supplementary Figure S3a). These activated Tregs (aTregs) were almost completely absent in the blood but were more than four times more abundant (p = 0.0001) in the tumor than in the peripheral tissue (Figure 3b). Furthermore, the expression of CD69 by Tregs in the tumor tissue was positively correlated with the mean expression of CTLA-4, TIGIT, ICOS, and PD-1 in CD8 effector memory cells, indicating a connection between immune checkpoint expression in effector cells and the activation status of regulatory T cells (Figure 3c). In addition to the well-studied CD4 Tregs, there was also an increase in CD8+ FoxP3+ CD25+ cells in the TILs population (Figure 3d). Although this population is considerably smaller, it is thought to have regulatory functions similar to those of CD4 Tregs and is more frequent in some types of cancers(31, 32).

### Lymph node-negative patients demonstrate enrichment of CD4 EM cells lacking multiple checkpoint co-expression

To identify the aggressive biology of the disease, we looked at immune cell distribution in lymph node-negative patients. We observed a cluster of CD4 EM cells enriched in the peripheral liver tissue of patients without lymph node invasion (Figure 3e). These CD4 EM cells are characterized by a lack of the expression of checkpoints CTLA-4, PD-1, and TIGIT as well as the activation marker CD69 (Supplementary Figure S3b). There was no correlation with overall survival, most likely due to the small cohort size and high postoperative mortality rate of almost 25%.

### PD-1 positive CD4 and CD8 residential effector memory cells are present in 24% of our patient cohort

PD-1 expression in effector memory cells is a potential biomarker to allow patient stratification for PD-1 blockade therapy. We applied multiplexed immunohistochemistry (IHC) on whole slide images to identify the spatial distribution of CD4 and CD8 T cells including CD103 as a marker for tissue-resident memory cells (TRMs), combined with PD-1 expression. CD4 and CD8 cells were mainly present in the tumor and interface areas, with and without PD-1 co-expression. The regions annotated by the pathologist as peripheral liver tissue showed lower levels of all the examined CD4 subsets (Figure 4a-j). Levels of PD-1 positive CD8 cells and CD8 TRMs were also more abundant in the tumor (T) and interface (I) regions compared to the peripheral tissue (P) (p = 0.003 (I-P) for CD8_PD-1 and p=0.05 (T-P), p=0.05 for CD8 TRM_PD1 (I-P) and p=0.02 (T-P)) (Figure 4f-j). These findings are in line with our CyTOF data, which demonstrated increased PD-1 expression in CD4 and CD8 TRMs in the tumor tissue. Also, CD4 and CD8 as well as PD-1 positive CD4 and CD8 cells were more abundant in the interface compared to the tumor region (p=0.02 for CD4, p=0.003 for CD8, p= 0.02 for CD4_PD-1, and p=0.002 CD8_PD-1). In our cohort, 24% of the patients (6 of 25 patients) demonstrated either CD4 or CD8 TRM cells being positive for PD-1, identifying a potential subgroup of patients who could respond to PD-1 blockade therapy. Of course, this finding would need further validation before this combination of markers could be used in the clinical field. Multiplexed immunohistochemistry is available in most university hospitals and could be a useful tool in the future since the spatiality of the immune cells is kept and plays a role.

**Figure 4.**
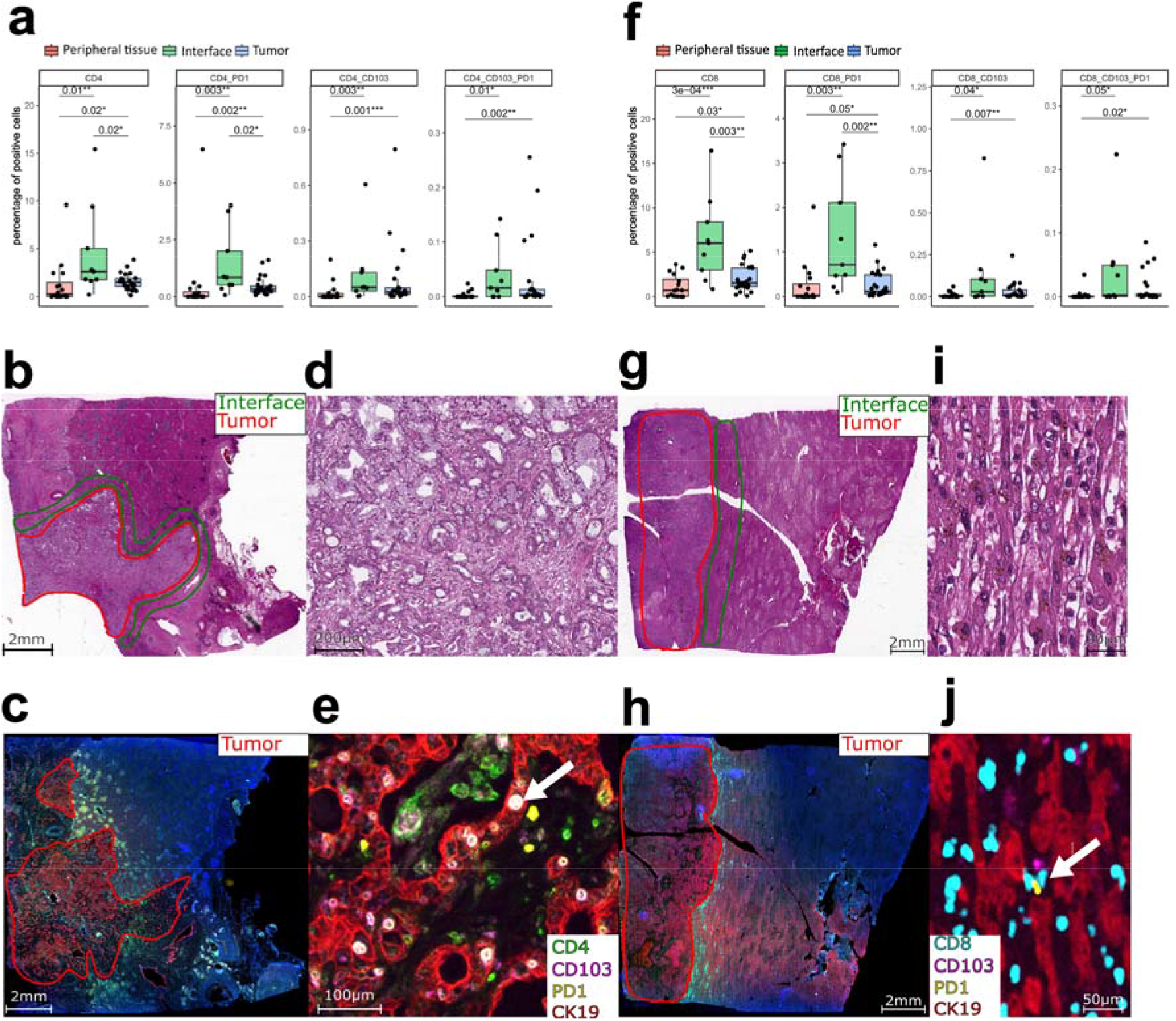
Validation by multiplexed immunohistochemistry. **(a)** Percentage of cells positive for CD4, CD103, and PD-1 in the different regions of interest (ROIs). PD-1/ CD103 positive cells were mainly found in the interface and tumor area, while positive cells for all markers were rare in the periphery **(b)** Overview of the HE-stained slide of patient 323. **(c)** The corresponding multiplexed image. **(d)** Magnified image of the HE-stained slide of the tumor area of patient 323. **(e)** The corresponding magnified multiplexed image. The highlighted cells with a box are CD4 effector memory cells with expression of PD-1. **(f)** Percentage of cells positive for CD8, CD103, and PD-1 in the different ROIs. Cells positive for the different markers were more abundant in the tumor and interface ROIs. **(g)** Overview of the HE-stained slide of patient 302. **(h)** The corresponding multiplexed image. **(i)** Magnified image of the HE-stained slide of the interface area of patient 302. **(j)** The corresponding magnified multiplexed image. The highlighted cells with a box are CD8 effector memory cells with expression of PD-1.

### Transcriptomics reveals heterogeneity in cholangiocarcinoma cancer cell populations

To understand cellular interactions between different cell types we applied transcriptomics. We performed snRNA sequencing of tumor tissue from seven patients with pCCA, split into two groups based on their clinical prognosis. Patients with cancer-positive lymph nodes and cancer recurrence within one year after surgery were identified as having a poor prognosis based on the aggressive biology of the disease (n=3 with good clinical prognosis; n=4 with poor clinical prognosis). Detailed patient characteristics are outlined in Supplementary Table S3 and available detected genetic variants are outlined in Supplementary Table S4. The tumor heterogeneity in iCCA is more diverse and for this reason, we chose pCCA samples.

We identified 10 distinct clusters of cells (Figure 5a), distributed across the two prognosis groups. These clusters were manually annotated into known cell types based on marker gene expression. A large proportion of cells, represented by 4 of the 10 identified clusters, expressed markers associated with cholangiocytes and malignant epithelial cells. The cholangiocyte cell cluster represent the cancer cells. Other identified cell clusters included T cells, B cells, and a cluster of macrophages and other myeloid cells as three distinct immune cell populations. Two clusters of hepatocytes and a cluster of stromal cells were identified (Supplementary Figure S4a). The different clusters were heterogeneously distributed between the patient and prognosis groups (Figure 5b). Hepatocytes were mainly present in only two patients, and the proportion of cancer cell clusters also differed between patients (Supplementary Figure S4b and S4c). For the total cancer cell population, we identified 4167 differentially upregulated and 5578 differentially downregulated genes (FDR<0.05) between prognosis groups, including MDM2 and PTPRM (Figure 5c and d).

**Figure 5.**
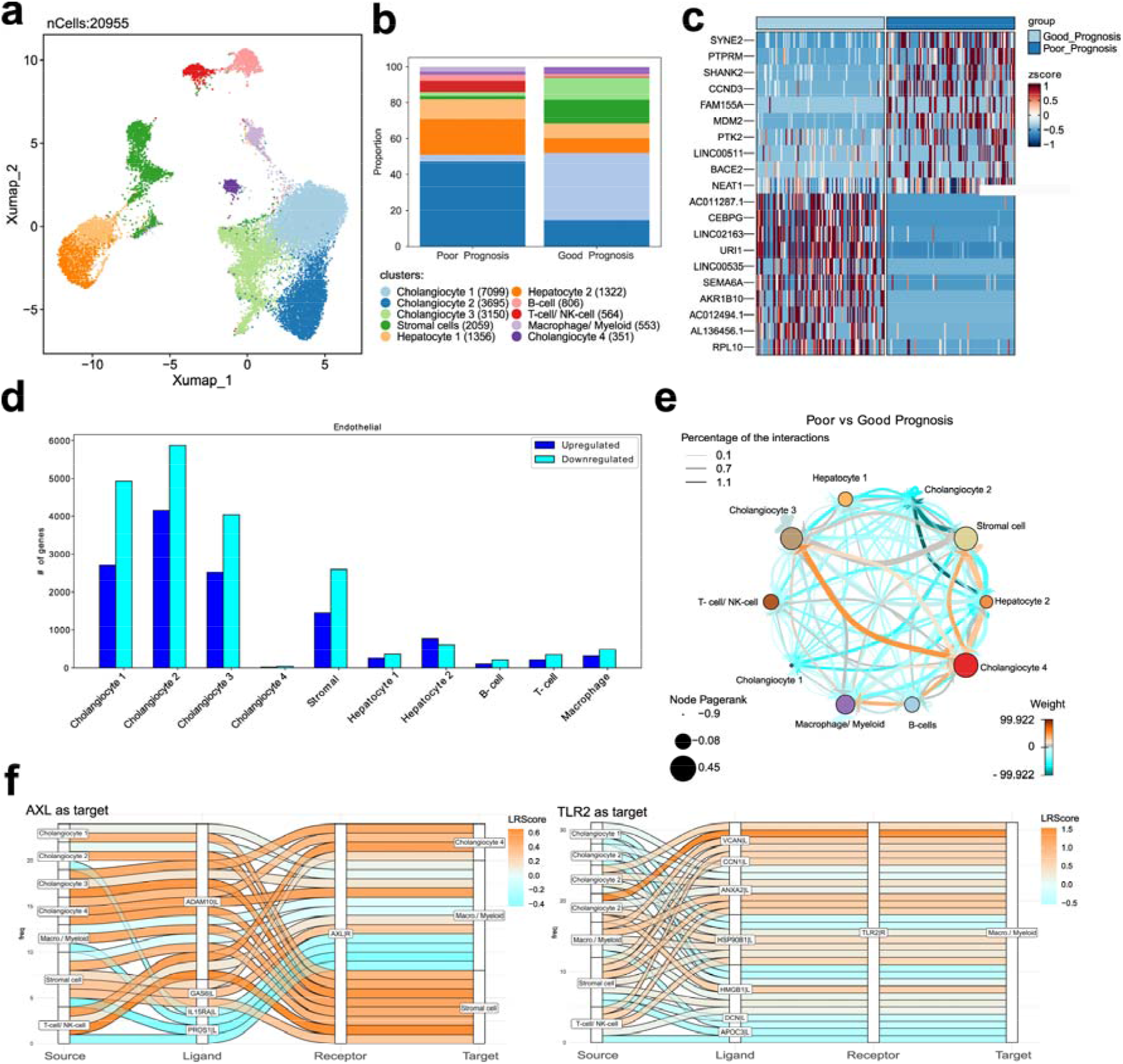
SnRNA reveals heterogeneity in cell clusters between clinical groups. **(a)** UMAP representation of clusters in single-cell data from 7 pCCA patients. **(b)** Stacked bar plot representing the compositional distribution of different cell types in good and poor prognosis. The cholangiocyte cell clusters represent the cancer cells. **(c)** Heat map plot representing top 10 differentially expressed up and down-regulated genes in poor prognosis taking good prognosis as a reference and all cancer cells taken together. **(d)** Bar plot showing the number of significantly up and down-regulated genes in different cell types in poor prognosis. **(e)** Network graph depicting comparative cell-cell communication based on good and poor prognosis, between different cell types. Here, the good prognosis is taken as a reference, and higher weight defines up-regulated communication in disease as compared to control. **(f)** Sankey plots visualize the top interactions of the target genes AXL (left) and TLR2 (right) predicted by the cell-cell communication analysis. Shown are Ligand receptor pairs as well as the cell types expressing the ligand (source) and receptor (target).

### Poor prognosis patients demonstrate an upregulation in crosstalk between macrophages, stromal cells, and cancer cells

Based on ligand-receptor expression, we assessed the cellular crosstalk between different cell populations. For this analysis, we used the CellCall(33) tool and uncovered the cell-cell communication between stromal cells, cancer cells, macrophages, and myeloid cells in both prognosis groups separately. Next, we applied the crosstalker tool to directly compare changes in cell-cell communication between the two prognosis groups. We observed a shift in cellular communication towards cancer cell cluster 4 in the poor prognosis group. This is mainly driven by communication between cancer cell cluster 4 and macrophages and myeloid cells, as well as the stromal cell cluster (Figure 5e and Supplementary Figure S4d). The top ligand in the crosstalk from cancer cell cluster 4 to the stromal and macrophage myeloid cell clusters was the Versican gene, VCAN. VCAN encodes an extracellular matrix protein known to be involved in cancer progression and inflammation(35). VCAN was predicted to interact with Toll-like receptor 2 (TLR2) on macrophages (Figure 5f) and epithelial growth factor receptor (EGFR) in stromal cells (Supplementary Figure S4e). Among the top ligands driving communication from the macrophage myeloid cell cluster was the tumor growth factor β (TGF-β) gene (Supplementary Figure S4e), TGFB1. TGF-β has two receptors, type I and type II, and is a well-known regulator of cancer immune evasion as well as other aspects of tumor progression(36). We also found AXL among the ligand-receptor pairs mainly present in the poor prognosis patients. AXL is a member of the three known TAM receptors (Tyro3, MerTK, AXL) interacting with its ligand GAS6. Specifically, GAS6 – AXL interaction was predicted between GAS6-expressing stromal cells, cancer cells and AXL-expressing macrophages (Figure 5f). TAM receptors (TAMr) are well-known regulators of macrophage function and are potent inhibitors of the inflammatory response.

### Poor prognosis signature is in a large subset of eCCA patients in available transcriptomic data

To validate the transcriptomic findings in our pCCA cohort, we analyzed publicly available microarray data from 182 cases of extrahepatic CCA (eCCA). The dataset contains dCCA cases as well, but the majority of 130 cases consist of pCCA(37). To assess cell type abundance, we performed cellular deconvolution using CIBERSORTx(38) based on signatures generated from the 10 clusters identified in our snRNA seq data. These predicted fractions were used to cluster the patients into 3 groups (Supplementary Figure S5a and S5b). While cancer cell clusters 1-3, stromal cells, both hepatocyte clusters and the macrophage/ myeloid cell cluster were predicted to be abundant, cancer cell cluster 4, B-cell and T-cell/ NK-cell fractions were predicted to be low (Figure 6a). Cluster 3 including 58 patients (32%) was predicted to be particularly rich in macrophages and stromal cells (Figure 6b). The same cluster expresses high levels of VCAN, EGFR, GAS6, and AXL (Figure 6c) suggesting the involvement of the two ligand-receptor pairs we identified in the poor prognosis patients in our snRNA sequencing data.

**Figure 6.**
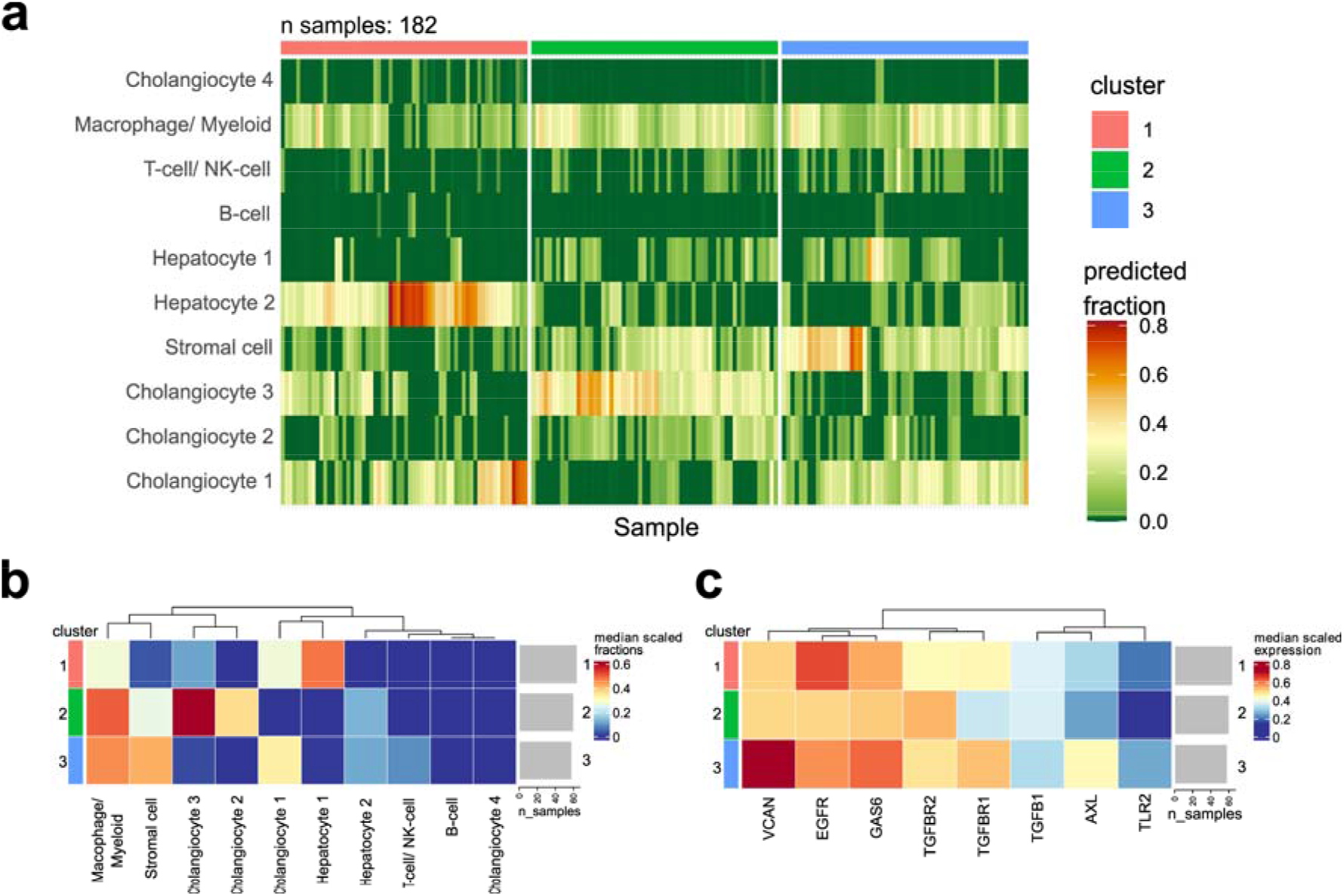
Poor prognosis signature is common in a subset of eCCA patients. Deconvolution of transcriptional data of 184 eCCA samples based on cibersortx using signatures generated from the clusters in our snRNA data. **(a)** Relative fraction predicted by cibersort for each sample. Samples were clustered into 3 distinct clusters based on the relative fractions. **(b)** Median relative fractions for the identified clusters. Cluster 3 is enriched in stromal cells and has a high abundance of macrophage/myeloid cells. **(c)** Gene expression of genes, identified to be involved in the cellular crosstalk in the TME of patients with a poor prognosis. Cluster 3 has high expression of VCAN-EGFR and GAS6-AXL ligand-receptor pairs.

## Discussion

CCA treatment options remain limited, and even after decades of research, surgery still represents the best chance for a cure. While the role of immunotherapy is increasing in other cancer entities, knowledge about immune responses to cancer cells in CCA remains limited. Furthermore, studies are complicated by CCA heterogeneity, including a wide range of genetic subtypes(39). The immune system can destroy cancer cells, but there is also a complicated immune activation network regulated by immune checkpoints(40). The TME of CCA features a T-cell exclusion mechanism and is dominated by low cytotoxic immune cell infiltration and high levels of Tregs and checkpoint expression, which hinders the immune response against tumors(5). Here, we report the contributions of different immune cells to the tumor microenvironment in iCCA and pCCA, with detailed definitions of their components.

Overall, the presence of many CD4 and CD8 cells in the tumor area is a sign of good prognosis, and previous studies have shown higher levels of CD4 and CD8 cells in the tumor margin than in the central tumor area in patients with iCCA(41, 42). Accumulation of Tregs in tumors correlates with poor prognosis(9, 43). Here, we confirmed previous findings, both iCCA and pCCA samples were infiltrated with Tregs and exhausted CD4 and CD8 tissue-resident memory cells, while CD8 effector memory cells were mostly excluded. Moreover, we identified elevated levels of CD69 in infiltrating Tregs, making them activated Tregs. Patients with activated Tregs, also demonstrated other immune cells with multiple co-expression of checkpoints, indicating an exhausted phenotype. CD69, as an activation marker, is involved in Treg function and has been reported to be correlated with tumor progression in hepatocellular carcinoma(44, 45).

Immune checkpoints play a role in many cancer types and only a subgroup of patients is likely to benefit from immunotherapy. An extensive understanding of immune cell composition and expression of costimulatory and coinhibitory checkpoints is needed to predict whether patients will respond to immunotherapy. PD-1 is a co-inhibitory receptor that can be expressed in many different cell types, including Tregs, activated B cells, and T cells. It suppresses the activity of T cells and downregulates the T cell response when it binds to its ligand PD-L1(19, 46). Our data showed high levels of PD-1 as well as other checkpoint inhibitors such as TIGIT and CTLA-4, confirming a dominant exhausted phenotype. Interestingly, in 24% of our patients, we identified CD4 and CD8 effector memory cells with co-expression of PD-1, this indicates a group of patients who could potentially benefit from immune blockade therapy. Further research is needed to validate this finding, but multiplexed immunohistochemistry seems to be a reliable tool to identify relevant immune cells while maintaining the spatial architecture.

Our transcriptomic data revealed the TME in a broader perspective and identified the crosstalk between cell clusters responsible for the immunosuppressive environment. We identified several immune populations, as well as clusters of stromal and cancer cells. Markers associated with stellate cells, which are believed to be a major source of CAFs(9, 47) were also present in our cohort. Following previous studies, we demonstrated increased expression of genes associated with a worse prognosis in the cancer cells, including MDM2 and PTPRM. MDM2 has been associated with advanced CCA stages and a worse clinical outcome(48, 49). PTPRM is associated with a worse outcome in cervical cancer(50). Macrophages and myeloid cells were identified within the TME and are known to contribute to immunosuppressive cell crosstalk within iCCA(51).

In our pCCA transcriptomic data, we identified GAS6, and AXL as unique ligand-receptor pairs associated with poor prognosis. *GAS6* codes for the growth arrest-specific 6 protein, which can be recognized by the AXL receptor belonging to the TAM subfamily. TAM receptors (TAMr) play a role in the immune system but are also associated with tumor progression(52). Both GAS6 and AXL are associated with worse patient survival in lung cancer(53), and their signaling is associated with the recruitment and polarization of suppressive macrophages in hepatocellular carcinoma(54). In addition to its immunosuppressive role, AXL and TAMr signaling are generally associated with PD-L1 expression(55, 56). Furthermore, we observed a shift in the crosstalk between different cell clusters. In our poor prognosis group, we identified increased crosstalk between a cancer cell subpopulation, macrophages, and stromal cells. Communication with macrophages was partly driven by the *VCAN* gene encoding for the extracellular matrix (ECM) protein versican and toll-like receptor 2 (TLR2). This interaction has been identified to promote an immunosuppressive environment(57) and to stimulate metastasis in cancer(58). In addition, versican is predicted to interact with the epithelial growth factor receptor (EGFR) in stromal cells. Versican signaling via EGFR in breast cancer has been shown to promote proliferation(59). EGFR is commonly overexpressed in CCA, which can be associated with unfavorable patient outcomes(60). We also identified the TGF-β gene as a major ligand expressed in macrophages. TAM-derived TGF-β is known to promote tumor progression(61) and drive the epithelial-mesenchymal transition in hepatocellular carcinomas(62). TGF-β is also a key mediator in the induction of Tregs and can directly inhibit cytotoxic T cell function(26, 63), making it a potent promoter for the suppression of T cells. In our poor prognosis group, we identified several ligand-receptor pairs, that are important mediators in the immune exhaustion of CCA and explain the limited immune cell-mediated anti-tumor response in this cancer entity.

In summary, we used high-dimensional phenotyping, transcriptomics, and multiplexed immunohistochemistry to provide a deeper insight into the immune exhausted phenotype in CCA. The multiple mechanisms leading to the exclusion of relevant immune cells needed for an anti-cancer response and mechanisms leading to active immune suppression are part of complex cell-cell crosstalk. Our findings confirmed the presence of Tregs and the lack of effector memory cells in the tumor.

New findings are the spatiality of the effector memory cells being more present in the peripheral tissue, for some reason these immune cells fail to reach the tumor niche.

Transcriptomic data identified the presence of specific cancer cells, mainly only present in the poor prognosis group. Here, we identified an upregulated crosstalk of these cancer cells with the myeloid compartment and the stromal cells. This crosstalk creates an immunosuppressive TME, worsening patient outcomes. Amongst the responsible ligand pairs are GAS6-AXL belonging to the TAM family. We then identified VCAN-TLR2 to be present and influencing the ECM in a way to supports immune exhaustion. Last, EGFR-TGF-β is expressed in macrophages and this finding is important in Tregs induction and blocking cytotoxic T cell function. Using deconvolution of previously published transcriptomic data in eCCA, we identified the involved cell types as well as VCAN, EGFR, GAS6, and AXL expression to be common in a subgroup of patients. This represents the poor prognostic group of patients with a more aggressive biology of the disease.

The immunosuppressive environment in CCA is a complex problem that needs to be tackled in the hope of creating better perspectives and therapy options for patients with CCA. In this cancer entity, only a limited response to checkpoint blockade has been observed. This study contributes to the understanding of the immunosuppressive environment and provides signaling pathways as drivers. Further examination of these pathways will provide valuable insights into the underlying mechanisms and could potentially give rise to treatment candidates that are tailored to the TME within CCA.

Good biomarkers for predicting a response to immune blockade treatment are still missing. PD-L1 immunohistochemistry is currently used in daily diagnostic care, and its expression in macrophages, immune cells, and cancer cells has been previously reported. Our study demonstrates that in addition to the presence of PD-L1 blocking immune cells in the tumor, the presence of immune cells outside the tumor in the peripheral surrounding liver tissue might be of importance for patient prognosis. A multiplexed panel can be used on peripheral liver biopsies to define the presence of CD4 and CD8 effector memory cells to predict an activated immune phenotype. We hypothesize that further research should investigate peripheral liver tissue for the presence of immune cell clusters and potentially identify response biomarkers for immunotherapy treatment. Although this is a relatively complex workup, multiplexed imaging is available in most university hospitals and may represent a step towards personalized cancer care in a patient group with sparse therapeutical options.

## Materials and Methods

### Patient Recruitment

Samples were collected between May 2020 and June 2021 from 16 patients with iCCA (n=7) or pCCA (n=9) with a curative intent for surgery. The patient characteristics are shown in Supplementary Table S1. All patients consented to participate in this study with local ethical approval under EK106/18 and EK360/19 and the broad consent (EK206/09) of the hospitals centralized biomaterial bank (RWTH cBMB).

### Sample Collection and Tissue Digestion

Blood and tissue samples were collected from each patient. Blood was collected before the operation in 20 ml EDTA tubes. Blood samples were processed within 4h of blood withdrawal. PBMCs were isolated using Ficoll after the addition of 20 mL of 2% FBS/PBS. After centrifuging for 20 minutes at 1300 g without break, we removed the PBMC ring using a pipette. This was then washed and centrifuged again at 300 g and if red blood cells were lysed with ACK lysing buffer the cells were washed again.

Tissue samples were collected postoperatively from the pathology department of our hospital. Tissue blocks of normal tissue and tumor tissue with a size of 10 × 10 × 5 mm were placed in RPMI medium (Gibco), which was cooled on ice. Our digestion protocol was used as in our previous study(64). The tissue was minced with a scalpel, and enzymes were added for digestion. This solution, along with the tissue pieces, was placed in a shaker for 40 min at 37 degrees Celsius (°C). Then, the medium was refreshed, new enzymes were added and again placed into the shaker for an additional 30 min. The remaining solid tissue components were pushed through using a cell strainer, providing the greatest possible yield. The tube was centrifuged at 300 g for 10 min, and the pellet was collected and cryopreserved in a medium containing DMSO at -80°C until further analysis.

### CyTOF sample preparation

The metal-isotope labeled antibodies used in this study were conjugated using the MaxPar X8 Antibody labeling kit according to the manufacturer’s instructions (Fluidigm) or were purchased from Fluidigm pre-conjugated. Each conjugated antibody was quality-checked and titrated to the optimal staining concentration using a combination of primary human cells (peripheral blood mononuclear cells, single-cell suspensions from tonsils, and single-cell suspension from cholangiocarcinoma). The samples were thawed for further CyTOF staining by gently diluting DMSO in warm RPMI and washing 2 times in RPMI. Single-cell suspensions of cholangiocarcinoma tumor tissue, surrounding normal liver tissue, and peripheral blood mononuclear cells were thawed and washed with warm culture media (RPMI + 10%FCS Pen/strep/glut). The cells were incubated with culture medium + 15 U/ml DNase for 10 min at 37°C. Cells were washed in PBS and incubated with 0.5 μM cisplatin (Fluidigm) in phosphate-buffered saline (PBS) for 5 min at room temperature (RT) to label non-viable cells. The cells were resuspended in Truestain FC blocker (BioLegend) for 10 min at RT. After incubation, an antibody mix of targets not capable of fixation (CCR5, CXCR3, CCR, and IL-7RA) was added to the cells for 30 min at 4°C. The cells were fixed with 1.6% PFA in PBS for 10 min at RT. Cells were washed with CSM and fixed with 1.6% PFA in PBS for 10 min at room temperature (RT). Fixed cells were palladium barcoded (Fluidigm) according to established protocols. All surface staining was performed in CSM for 30 min at RT (antibodies listed in Supplementary Table S2). Cells were washed in CSM and permeabilized using the FoxP3 staining kit, according to the manufacturer’s instructions (eBioscience™ FoxP3/Transcription Factor Staining Buffer Set, Invitrogen. After intracellular staining, cells were fixed with 1.6% PFA in PBS for 10 min at RT. Before acquisition, samples were washed in CSM and resuspended in intercalation solution (1.6% PFA in PBS, 0.02% saponin (Sigma), and 0.5 μM iridium diluted in Fix/Perm solution (Fluidigm)) for 1 h at RT or overnight at 4°C or frozen in aliquots to label DNA. Before acquisition, the cells were washed once with RPMI+30% FCS. The samples were washed twice with CSB and once with larger volumes of CAS. All samples were filtered through a 35 μM nylon mesh cell strainer, resuspended at 0,5 × 106 cells/mL in CAS supplemented with 1/10 EQ four-element calibration beads (Fluidigm), and acquired on a CyTOF2 mass cytometer (Fluidigm) using the Super Sampler injection system. Before data acquisition, the instrument was demonstrated to have Tb159 dual counts greater than 1,000,000 and oxidation less than 3%; if the instrument failed these criteria, it was cleaned, tuned, or repaired as necessary.

### CyTOF data acquisition and pre-analysis

Immediately before the acquisition, samples were washed as described above and then resuspended at a concentration of 0.5 million cells/mL in cell acquisition solution containing a 1/10 dilution of EQ 4 Element Beads (Fluidigm). Samples were acquired on a CyTOF Helios mass cytometer at an event rate of <150 events/s. After the acquisition, the data were normalized using bead-based normalization in CyTOF software. The data were exported as FCS files for downstream analyses. The data were gated to exclude residual normalization beads, debris, dead cells, and doublets, leaving DNA+ live single events for subsequent clustering and high-dimensional analysis.

### Batch effect correction

Data were acquired in five different batches, each containing two technical replicates, one single-cell suspension of tonsil cells, and one PBMC sample. We used CytoNorm(65) to model the differences in signal intensities between batches. We determined which of the two replicate sets had a larger dynamic range per marker. TIGIT, CD69, CTLA-4, ICOS, PD-1, CXCR3, LAG-3, OX40, CD7, CD95, CD45RA, CD45, CD127, and HLA-DR were modeled and normalized using tonsil replicates. All other markers were modeled using PBMC replicates. The data were arcsinh-transformed (factor 5) and clustered into B cells, T cells, NK cells, and myeloid lineages using FlowSOM. Subsequent settings for CytoNorm were as follows: 21 quantiles from the 0th to 99th percentile. The limits were (0 and 7.4). All FCS files were normalized using both normalization models.

### CyTOF data analysis

After batch correction was performed, the data were further analyzed using R (v. 4.2.2). Subsequently, cells were clustered using the FlowSOM(66) algorithm (CATLYST package v. 1.22.0) based on CD2, CD3, CD4, CD8, CD14, CD45, and HLA-DR on a 10 by 10 grid and with elbow metaclustering. The algorithm resulted in eight metaclusters that could be manually assigned to CD4 T cells (FlowSOM metaclusters 1 and 4), CD8 T cells (FlowSOM metaclusters 2 and 3), and miscellaneous cells, included NK cells, CD14+ cells, and others (FlowSOM metaclusters 5-8).

Cells were downsampled to less than 50000 CD4 and CD8 cells per sample, resulting in 918.019 CD4 T cells and 732.047 CD8 T cells. OptSNE dimensionality reduction from the MulticoreTSNE package (v.0.1) using Python (v.2.7.18) with a perplexity of 30, theta of 0.5, maximum of 1000 iterations, and stop of 5000 was performed using CD45RA, CD45RO, CCR7, CD27, CD28, CD69, CD103, CD95, CD57, CD161, CD44, CD25, CD127, FOXP3, CXCR3, CCR5, CCR4, CD11a, and CD49d. Subsequently, Phenograph(67) (Rphenograph package v. 0.99.1) was applied using the same markers with K-nearest neighbors set to 20 and Euclidean as the distance metric. The resulting clusters (25 for CD4 + T cells and 31 for CD8 + T cells) were annotated according to their expression profiles into known immune cell subset categories for data interpretation.

Statistical comparisons involved Friedman’s test to compare mean frequencies and marker expression across tissues in either iCCA or pCCA (matched blood, normal tissue, and tumor tissue samples), Mann-Whitney test to compare mean frequencies and marker expression between iCCA and pCCA tumors, and Wilcoxon test to compare mean frequencies and marker expression between normal and tumor tissues. Correlation analysis between cluster frequencies in blood versus tumor tissues and mean marker expression was performed using Spearman’s test.

### Histology and multiplexed immunohistochemistry

The histology report of each patient was checked for quality control of the right tumor. The pathologist (LH) collected the H&E-stained slides and selected the tumor block. Sections were cut using a Leica RM2235 microscope at a thickness of approximately 4 μm. Antibodies against CK19, CD4, CD8, CD103, and PD-1 were analyzed, and their co-expression was quantified using the TissueFAXS PLUS scanning system with StrataQuest Analysis Software from TissueGnostics(68), Vienna, Austria.

#### Single-nuclei isolation

In addition to cytometric analysis, we also chose seven patients for transcriptomic analysis using single nuclei RNA sequencing (snRNA seq). The patients were divided into two groups based on their clinical prognosis. For the four patients in the poor clinical prognosis, we detected lymph node invasion by the tumor and recurrence within 1 year of surgery. In the three patients in the good prognosis group, no lymph node invasion was detected, and they were recurrence-free at the time of the study (at least one year after surgery). NGS data was available in 3 patients, see Supplementary Table S4. Tumor samples were frozen in cryotubes without fluid as quickly as possible after sample collection. To do so, they were covered in liquid nitrogen and then transferred to a −80□°C freezer for long-term storage. Single-nuclei isolation was performed as described previously(69).

Briefly, tissue samples were cut into pieces <0.5□cm and homogenized using a glass Dounce tissue grinder (Sigma-Aldrich). The tissue was homogenized by subjecting it to 25 strokes with pestle A and 25 strokes with pestle B in 2□ml of ice-cold nuclei EZ lysis buffer. Subsequently, the cells were incubated with an additional 3□ml of cold EZ lysis buffer on ice for 5□min. Nuclei were centrifuged at 500 g for 5□min at 4□°C, washed with 5□ml ice-cold EZ lysis buffer, and incubated on ice for 5□min. After centrifugation, the resulting nuclei pellet was washed with 5□ml nuclei suspension buffer (NSB; consisting of 1× PBS, 0.01% BSA, and 0.1% RNase inhibitor). Isolated nuclei were resuspended in 2□ml NSB, filtered through a 35□μm cell strainer, and counted. Approximately 1,000□nuclei per µL were used for loading on a 10x channel.

#### snRNA-seq preprocessing, clustering, and annotation

From the single nuclei using 10X sequencing, we generated raw reads that were further processed by Cellranger version 6.1.2, using GRCh38-2020-A as the human reference transcriptome. This gave us a count matrix that was converted to a fast q file by the default mkfastq function, followed by alignment using the count function, cell calling, and quantification of UMI counts. Only cells that expressed between 1000 and 20000 genes were retained using the filter_cells function. Furthermore, cells with more than 10% of the reads mapped to the mitochondrial genes were removed. To exclude potential doublets, the Scrublet tool was used, with a cutoff of 0.2. For subsequent analysis, we used Seurat (v. 4.3.0). The expression data were log-normalized with a scaling factor of 10000. We applied the Seurat V3 method to select the top 2000 highly variable genes that were subsequently used to calculate the principal components of the data. The harmony package was utilized as a batch-effect correction tool and set on sample IDs as the batch variable. UMAP was applied with Seurat’s RunUMAP function using the first 30 principal components generated from the harmony corrected PCA embedding. The cells were clustered using the Louvain algorithm at resolutions ranging from 0.3 1.1. To determine the final resolution, we manually inspected the marker genes and cluster composition. The dataset used for further downstream applications consisted of 20995 cells from 7 human CCA samples. Clusters were annotated using curated marker genes from literature.

#### Differential expression analyses

We used the MAST algorithm to identify differentially expressed genes in all cell types based on the prognosis groups. First, genes with low expression and normalized expression below zero were removed. The hurdle model was set with the good prognosis group as the reference condition and the poor prognosis as our condition. We extracted the differentially expressed genes, with the logarithm of fold change (logFC) value and adjusted p values from the summary statistics of the output. The Benjamini-Hochberg FDR correction method was used to identify statistically significant DE genes. Gene set enrichment analysis (GSEA) of differentially expressed genes was performed using the clusterProfiler package (v. 3.17) with a minimal gene set size of 3, a maximal gene set size of 500, and a p-value cutoff of 0.05 using Gene Ontology annotations.

#### Cell-cell communication analyses

To predict ligand receptor (LR) interactions between the previously identified clusters, we used the CrossTalkeR package (v. 1.3.6). The *liana_wrap* function of liana (v. 0.1.12) was used to generate LR prediction. As an input, CrossTalkeR uses cellphoneDB predictions as a database and generates a comparative cell-cell interaction (CCI) network for the conditions. After our unbiased analysis of LR all pairs, we focused on pairs that were among the top 10 pairs with the highest LR scores within the identified cell clusters in the CCI network using the prognosis groups as conditions. We also used the CellCall package (v.1.0.7) to identify cell-to-cell communication trends along different clusters, and internal regulatory signals. The *TransCommuProfile* function in the CellCall package determines the LR score between cell types. We used default parameters (0.05) for pvalue, Cor Spearson correlation significance and p.adjust value for regulon’s GSEA. The cellcall tool uses LR information from the KEGG database to infer the communication and transcription factor activity along with analyzing crucial pathway activity involved in communication between certain cellular connections.

### Analysis of public expression data

Transcriptome array data were retrieved from the Gene Expression Omnibus (GEO) dataset GSE132305 for 182 patients with eCCA, in this cohort 130 cases of pCCA were included and processed according to the authors’ original methodology (37). For cellular deconvolution, expression data were exported and analyzed using the CIBERSTORx tool(38). The signature matrix file was generated using the snRNA and clusters from this study as a single cell reference, as described in the original article. For the signature matrix, we used the top 50 DE genes from each of the 10 clusters resulting in 337 unique genes. The estimated relative fractions were used as input for clustering via phenograph.

## References

1. Bertuccio P, Malvezzi M, Carioli G, Hashim D, Boffetta P, El-Serag HB, et al. Global trends in mortality from intrahepatic and extrahepatic cholangiocarcinoma. J Hepatol. 2019;71(1):104–14.

2. Banales JM, Marin JJG, Lamarca A, Rodrigues PM, Khan SA, Roberts LR, et al. Cholangiocarcinoma 2020: the next horizon in mechanisms and management. Nat Rev Gastroenterol Hepatol. 2020;17(9):557-88.

3. Goeppert B. [Cholangiocarcinoma-diagnosis, classification, and molecular alterations]. Pathologe. 2020;41(5):488–94.

4. Bagante F, Spolverato G, Weiss M, Alexandrescu S, Marques HP, Aldrighetti L, et al. Assessment of the Lymph Node Status in Patients Undergoing Liver Resection for Intrahepatic Cholangiocarcinoma: the New Eighth Edition AJCC Staging System. J Gastrointest Surg. 2018;22(1):52–9.

5. Zhou G, Sprengers D, Mancham S, Erkens R, Boor PPC, van Beek AA, et al. Reduction of immunosuppressive tumor microenvironment in cholangiocarcinoma by ex vivo targeting immune checkpoint molecules. J Hepatol. 2019;71(4):753–62.

6. Bednarsch J, Tan X, Czigany Z, Liu D, Lang SA, Sivakumar S, et al. The Presence of Small Nerve Fibers in the Tumor Microenvironment as Predictive Biomarker of Oncological Outcome Following Partial Hepatectomy for Intrahepatic Cholangiocarcinoma. Cancers. 2021;13(15):3661.

7. Bednarsch J, Kather J, Tan X, Sivakumar S, Cacchi C, Wiltberger G, et al. Nerve Fibers in the Tumor Microenvironment as a Novel Biomarker for Oncological Outcome in Patients Undergoing Surgery for Perihilar Cholangiocarcinoma. Liver Cancer. 2021.

8. Loeuillard E, Conboy CB, Gores GJ, Rizvi S. Immunobiology of cholangiocarcinoma. JHEP reports: innovation in hepatology. 2019;1(4):297–311.

9. Fabris L, Sato K, Alpini G, Strazzabosco M. The Tumor Microenvironment in Cholangiocarcinoma Progression. Hepatology (Baltimore, Md). 2021;73 Suppl 1(Suppl 1):75-85.

10. Fabris L, Perugorria MJ, Mertens J, Björkström NK, Cramer T, Lleo A, et al. The tumour microenvironment and immune milieu of cholangiocarcinoma. Liver international: official journal of the International Association for the Study of the Liver. 2019;39 Suppl 1:63–78.

11. Alvisi G, Termanini A, Soldani C, Portale F, Carriero R, Pilipow K, et al. Multimodal single-cell profiling of intrahepatic cholangiocarcinoma defines hyperactivated Tregs as a potential therapeutic target. J Hepatol. 2022;77(5):1359–72.

12. Zhang M, Yang H, Wan L, Wang Z, Wang H, Ge C, et al. Single-cell transcriptomic architecture and intercellular crosstalk of human intrahepatic cholangiocarcinoma. J Hepatol. 2020;73(5):1118–30.

13. Clark CE, Hingorani SR, Mick R, Combs C, Tuveson DA, Vonderheide RH. Dynamics of the immune reaction to pancreatic cancer from inception to invasion. Cancer research. 2007;67(19):9518–27.

14. Kasper HU, Drebber U, Stippel DL, Dienes HP, Gillessen A. Liver tumor infiltrating lymphocytes: comparison of hepatocellular and cholangiolar carcinoma. World J Gastroenterol. 2009;15(40):5053–7.

15. Asahi Y, Hatanaka KC, Hatanaka Y, Kamiyama T, Orimo T, Shimada S, et al. Prognostic impact of CD8+ T cell distribution and its association with the HLA class I expression in intrahepatic cholangiocarcinoma. Surg Today. 2020;50(8):931–40.

16. Konduri V, Oyewole-Said D, Vazquez-Perez J, Weldon SA, Halpert MM, Levitt JM, et al. CD8(+)CD161(+) T-Cells: Cytotoxic Memory Cells With High Therapeutic Potential. Front Immunol. 2020;11:613204.

17. Li Z, Zheng B, Qiu X, Wu R, Wu T, Yang S, et al. The identification and functional analysis of CD8+PD-1+CD161+ T cells in hepatocellular carcinoma. NPJ Precis Oncol. 2020;4:28.

18. Zhou X, Du J, Liu C, Zeng H, Chen Y, Liu L, et al. A Pan-Cancer Analysis of CD161, a Potential New Immune Checkpoint. Front Immunol. 2021;12:688215.

19. Tian L, Ma J, Ma L, Zheng B, Liu L, Song D, et al. PD-1/PD-L1 expression profiles within intrahepatic cholangiocarcinoma predict clinical outcome. World journal of surgical oncology. 2020;18(1):303.

20. Ma L, Wang L, Khatib SA, Chang CW, Heinrich S, Dominguez DA, et al. Single-cell atlas of tumor cell evolution in response to therapy in hepatocellular carcinoma and intrahepatic cholangiocarcinoma. J Hepatol. 2021;75(6):1397–408.

21. Zhou M, Wang C, Lu S, Xu Y, Li Z, Jiang H, et al. Tumor-associated macrophages in cholangiocarcinoma: complex interplay and potential therapeutic target. EBioMedicine. 2021;67:103375.

22. Hasita H, Komohara Y, Okabe H, Masuda T, Ohnishi K, Lei XF, et al. Significance of alternatively activated macrophages in patients with intrahepatic cholangiocarcinoma. Cancer science. 2010;101(8):1913–9.

23. Atanasov G, Dietel C, Feldbrugge L, Benzing C, Krenzien F, Brandl A, et al. Tumor necrosis and infiltrating macrophages predict survival after curative resection for cholangiocarcinoma. Oncoimmunology. 2017;6(8):e1331806.

24. Sun D, Luo T, Dong P, Zhang N, Chen J, Zhang S, et al. CD86(+)/CD206(+) tumor-associated macrophages predict prognosis of patients with intrahepatic cholangiocarcinoma. PeerJ. 2020;8:e8458.

25. Loeuillard E, Yang J, Buckarma E, Wang J, Liu Y, Conboy C, et al. Targeting tumor-associated macrophages and granulocytic myeloid-derived suppressor cells augments PD-1 blockade in cholangiocarcinoma. J Clin Invest. 2020;130(10):5380–96.

26. Tauriello DVF, Palomo-Ponce S, Stork D, Berenguer-Llergo A, Badia-Ramentol J, Iglesias M, et al. TGFbeta drives immune evasion in genetically reconstituted colon cancer metastasis. Nature. 2018;554(7693):538-43.

27. Kalluri R. The biology and function of fibroblasts in cancer. Nat Rev Cancer. 2016;16(9):582–98.

28. Huang B, Liu R, Wang P, Yuan Z, Yang J, Xiong H, et al. CD8(+)CD57(+) T cells exhibit distinct features in human non-small cell lung cancer. J Immunother Cancer. 2020;8(1).

29. Strioga M, Pasukoniene V, Characiejus D. CD8+ CD28- and CD8+ CD57+ T cells and their role in health and disease. Immunology. 2011;134(1):17–32.

30. Ohue Y, Nishikawa H. Regulatory T (Treg) cells in cancer: Can Treg cells be a new therapeutic target? Cancer science. 2019;110(7):2080–9.

31. Flippe L, Bezie S, Anegon I, Guillonneau C. Future prospects for CD8(+) regulatory T cells in immune tolerance. Immunol Rev. 2019;292(1):209–24.

32. Chaput N, Louafi S, Bardier A, Charlotte F, Vaillant JC, Menegaux F, et al. Identification of CD8+CD25+Foxp3+ suppressive T cells in colorectal cancer tissue. Gut. 2009;58(4):520–9.

33. Zhang Y, Liu T, Hu X, Wang M, Wang J, Zou B, et al. CellCall: integrating paired ligand-receptor and transcription factor activities for cell-cell communication. Nucleic Acids Res. 2021;49(15):8520–34.

34. Lemke G, Rothlin CV. Immunobiology of the TAM receptors. Nat Rev Immunol. 2008;8(5):327–36.

35. Papadas A, Arauz G, Cicala A, Wiesner J, Asimakopoulos F. Versican and Versican-matrikines in Cancer Progression, Inflammation, and Immunity. J Histochem Cytochem. 2020;68(12):871–85.

36. Yang L, Pang Y, Moses HL. TGF-beta and immune cells: an important regulatory axis in the tumor microenvironment and progression. Trends Immunol. 2010;31(6):220–7.

37. Montal R, Sia D, Montironi C, Leow WQ, Esteban-Fabro R, Pinyol R, et al. Molecular classification and therapeutic targets in extrahepatic cholangiocarcinoma. J Hepatol. 2020;73(2):315–27.

38. Newman AM, Steen CB, Liu CL, Gentles AJ, Chaudhuri AA, Scherer F, et al. Determining cell type abundance and expression from bulk tissues with digital cytometry. Nat Biotechnol. 2019;37(7):773–82.

39. Rizvi S, Khan SA, Hallemeier CL, Kelley RK, Gores GJ. Cholangiocarcinoma - evolving concepts and therapeutic strategies. Nature reviews Clinical oncology. 2018;15(2):95–111.

40. Tumeh PC, Harview CL, Yearley JH, Shintaku IP, Taylor EJ, Robert L, et al. PD-1 blockade induces responses by inhibiting adaptive immune resistance. Nature. 2014;515(7528):568-71.

41. Carapeto F, Bozorgui B, Shroff RT, Chagani S, Solis Soto L, Foo WC, et al. The Immunogenomic Landscape of Resected Intrahepatic Cholangiocarcinoma. Hepatology (Baltimore, Md). 2021.

42. Goeppert B, Frauenschuh L, Zucknick M, Stenzinger A, Andrulis M, Klauschen F, et al. Prognostic impact of tumour-infiltrating immune cells on biliary tract cancer. Br J Cancer. 2013;109(10):2665–74.

43. Kitano Y, Okabe H, Yamashita YI, Nakagawa S, Saito Y, Umezaki N, et al. Tumour-infiltrating inflammatory and immune cells in patients with extrahepatic cholangiocarcinoma. Br J Cancer. 2018;118(2):171–80.

44. Zhu J, Feng A, Sun J, Jiang Z, Zhang G, Wang K, et al. Increased CD4(+) CD69(+) CD25(-) T cells in patients with hepatocellular carcinoma are associated with tumor progression. J Gastroenterol Hepatol. 2011;26(10):1519–26.

45. Gonzalez-Amaro R, Cortes JR, Sanchez-Madrid F, Martin P. Is CD69 an effective brake to control inflammatory diseases? Trends Mol Med. 2013;19(10):625–32.

46. Gutiérrez-Larrañaga M, González-López E, Roa-Bautista A, Rodrigues PM, Díaz-González Á, Banales JM, et al. Immune Checkpoint Inhibitors: The Emerging Cornerstone in Cholangiocarcinoma Therapy? Liver Cancer. 2021;10(6):545–60.

47. Okabe H, Beppu T, Hayashi H, Horino K, Masuda T, Komori H, et al. Hepatic stellate cells may relate to progression of intrahepatic cholangiocarcinoma. Ann Surg Oncol. 2009;16(9):2555–64.

48. Horie S, Endo K, Kawasaki H, Terada T. Overexpression of MDM2 protein in intrahepatic cholangiocarcinoma: relationship with p53 overexpression, Ki-67 labeling, and clinicopathological features. Virchows Archiv: an international journal of pathology. 2000;437(1):25-30.

49. Wattanawongdon W, Simawaranon Bartpho T, Tongtawee T. Expression of CD44 and MDM2 in cholangiocarcinoma is correlated with poor clinicopathologic characteristics. Int J Clin Exp Pathol. 2019;12(10):3961–7.

50. Liu P, Zhang C, Liao Y, Liu J, Huang J, Xia M, et al. High expression of PTPRM predicts poor prognosis and promotes tumor growth and lymph node metastasis in cervical cancer. Cell Death Dis. 2020;11(8):687.

51. O’Rourke CJ, Salati M, Rae C, Carpino G, Leslie H, Pea A, et al. Molecular portraits of patients with intrahepatic cholangiocarcinoma who diverge as rapid progressors or long survivors on chemotherapy. Gut. 2023.

52. Wu G, Ma Z, Hu W, Wang D, Gong B, Fan C, et al. Molecular insights of Gas6/TAM in cancer development and therapy. Cell Death Dis. 2017;8(3):e2700.

53. Ishikawa M, Sonobe M, Nakayama E, Kobayashi M, Kikuchi R, Kitamura J, et al. Higher expression of receptor tyrosine kinase Axl, and differential expression of its ligand, Gas6, predict poor survival in lung adenocarcinoma patients. Ann Surg Oncol. 2013;20 Suppl 3(Suppl 3):S467-76.

54. Yang F, Wei Y, Han D, Li Y, Shi S, Jiao D, et al. Interaction with CD68 and Regulation of GAS6 Expression by Endosialin in Fibroblasts Drives Recruitment and Polarization of Macrophages in Hepatocellular Carcinoma. Cancer research. 2020;80(18):3892–905.

55. Kasikara C, Kumar S, Kimani S, Tsou WI, Geng K, Davra V, et al. Phosphatidylserine Sensing by TAM Receptors Regulates AKT-Dependent Chemoresistance and PD-L1 Expression. Mol Cancer Res. 2017;15(6):753–64.

56. Tsukita Y, Fujino N, Miyauchi E, Saito R, Fujishima F, Itakura K, et al. Axl kinase drives immune checkpoint and chemokine signalling pathways in lung adenocarcinomas. Mol Cancer. 2019;18(1):24.

57. Hope C, Foulcer S, Jagodinsky J, Chen SX, Jensen JL, Patel S, et al. Immunoregulatory roles of versican proteolysis in the myeloma microenvironment. Blood. 2016;128(5):680–5.

58. Kim S, Takahashi H, Lin WW, Descargues P, Grivennikov S, Kim Y, et al. Carcinoma-produced factors activate myeloid cells through TLR2 to stimulate metastasis. Nature. 2009;457(7225):102-6.

59. Du WW, Fang L, Yang X, Sheng W, Yang BL, Seth A, et al. The role of versican in modulating breast cancer cell self-renewal. Mol Cancer Res. 2013;11(5):443–55.

60. Yoshikawa D, Ojima H, Iwasaki M, Hiraoka N, Kosuge T, Kasai S, et al. Clinicopathological and prognostic significance of EGFR, VEGF, and HER2 expression in cholangiocarcinoma. Br J Cancer. 2008;98(2):418-25.

61. Xiang X, Wang J, Lu D, Xu X. Targeting tumor-associated macrophages to synergize tumor immunotherapy. Signal Transduct Target Ther. 2021;6(1):75.

62. Fan QM, Jing YY, Yu GF, Kou XR, Ye F, Gao L, et al. Tumor-associated macrophages promote cancer stem cell-like properties via transforming growth factor-beta1-induced epithelial-mesenchymal transition in hepatocellular carcinoma. Cancer Lett. 2014;352(2):160–8.

63. Thomas DA, Massague J. TGF-beta directly targets cytotoxic T cell functions during tumor evasion of immune surveillance. Cancer Cell. 2005;8(5):369–80.

64. Sivakumar S, Abu-Shah E, Ahern DJ, Arbe-Barnes EH, Jainarayanan AK, Mangal N, et al. Activated Regulatory T-Cells, Dysfunctional and Senescent T-Cells Hinder the Immunity in Pancreatic Cancer. Cancers (Basel). 2021;13(8).

65. Van Gassen S, Gaudilliere B, Angst MS, Saeys Y, Aghaeepour N. CytoNorm: A Normalization Algorithm for Cytometry Data. Cytometry A. 2020;97(3):268–78.

66. Van Gassen S, Callebaut B, Van Helden MJ, Lambrecht BN, Demeester P, Dhaene T, et al. FlowSOM: Using self-organizing maps for visualization and interpretation of cytometry data. Cytometry A. 2015;87(7):636–45.

67. Levine JH, Simonds EF, Bendall SC, Davis KL, Amir el AD, Tadmor MD, et al. Data-Driven Phenotypic Dissection of AML Reveals Progenitor-like Cells that Correlate with Prognosis. Cell. 2015;162(1):184–97.

68. Tan X, Bednarsch J, Rosin M, Appinger S, Liu D, Wiltberger G, et al. PD-1+ T-Cells Correlate with Nerve Fiber Density as a Prognostic Biomarker in Patients with Resected Perihilar Cholangiocarcinoma. Cancers (Basel). 2022;14(9).

69. Habib N, Avraham-Davidi I, Basu A, Burks T, Shekhar K, Hofree M, et al. Massively parallel single-nucleus RNA-seq with DroNc-seq. Nat Methods. 2017;14(10):955–8.

