## Supplementary files for "Multimodal single-cell profiling reveals cancer crosstalk between macrophages and stromal cells in poor prognostic cholangiocarcinoma patients"

Table S1. Clinical data for patients included in the CyTOF cohort

| | iCCA | | pCCA | |
|---|---|---|---|---|
| **Included patients** | 6 | | 9 | |
| **Age in years** | 68.7 | | 64.2 | |
| **RFS in months** | 17.2 | | 11.8 | |
| **Survival time in months** | 22.8 | | 14.2 | |
| **Grade** | | | | |
| G1 | 0 | 0% | 0 | 0% |
| G2 | 3 | 50% | 1 | 11% |
| G3 | 2 | 33% | 5 | 56% |
| G4 | 1 | 17% | 3 | 33% |
| not known | 1 | 17% | 0 | 0% |
| **Perineural invasion** | | | | |
| positive | 2 | 33% | 1 | 11% |
| negative | 3 | 50% | 8 | 89% |
| not known | 1 | 17% | 0 | 0% |
| **Lymph node invasion** | | | | |
| positive | 4 | 67% | 3 | 33% |
| negative | 1 | 17% | 6 | 67% |
| not known | 1 | 17% | 0 | 0% |
| **Lymphovascular invasion** | | | | |
| positive | 0 | 0% | 2 | 22% |
| negative | 6 | 100% | 7 | 78% |
| not known | 0 | 0% | 0 | 0% |

Table S2. antibodies used for CyTOF

| Target protein | Clone | Metal | Manufacturer |
|---|---|---|---|
| **Barcodes** | | 103-110Pd | Standard BioTools |
| **Iridium** | | 191-193Ir | Standard BioTools |
| **Cisplatin** | | 194-195Pt | Standard BioTools |
| **CD45** | HI30 | 89Y | Standard BioTools |
| **CD49d** | 9F10 | 141Pr | Standard BioTools |
| **CD11a** | HI111 | 142Nd | Standard BioTools |
| **CD5** | UCHT2 | 143Nd | Standard BioTools |
| **CD195 (CCR5)** | NP-6G4 | 144Nd | Standard BioTools |
| **CD4** | RPA-T4 | 145Nd | Standard BioTools |
| **CD8a** | RPA-T8 | 146Nd | Standard BioTools |
| **CD7** | CD7-6B7 | 147Sm | Standard BioTools |
| **CD103** | Ber-ACT3 | 148Nd | BioLegend |
| **CD25 (IL-2R)** | 2A3 | 149Sm | Standard BioTools |
| **CD134 (OX40)** | ACT35 | 150Nd | Standard BioTools |
| **CD2** | TS1/8 | 151Eu | Standard BioTools |
| **CD95 (Fas)** | DX2 | 152Sm | Standard BioTools |
| **CD366 (TIM-3)** | F382E2 | 153Eu | BioLegend |
| **CD14** | 61D3 | 154Sm | Hybridoma |
| **CD279 (PD-1)** | EH12.2H7 | 155Gd | Standard BioTools |
| **CD183 (CXCR3)** | G025H7 | 156Gd | Standard BioTools |
| **CD194 (CCR4)** | L291H4 | 158Gd | Standard BioTools |
| **CD197 (CCR7)** | G043H7 | 159Tb | Standard BioTools |
| **CD28** | CD28.2 | 160Gd | Standard BioTools |
| **CD152 (CTLA-4)** | 14D3 | 161Dy | Standard BioTools |
| **CD69** | FN50 | 162Dy | Standard BioTools |
| **TIGIT** | MBSA43 | 163Dy | ThermoFisher |
| **CD161** | HP3G10 | 164Dy | Standard BioTools |
| **CD45RO** | UCHL1 | 165Ho | Standard BioTools |
| **CD44** | BJ18 | 166Er | Standard BioTools |
| **CD27** | 323 | 167Er | Standard BioTools |
| **CD278 (ICOS)** | C398.4A | 168Er | Standard BioTools |
| **CD45RA** | HI100 | 169Tm | Standard BioTools |
| **CD3** | UCHT1 | 170Er | Fluidigm |
| **GITR** | 621 | 171Yb | BioLegend |
| **CD57** | HCD57 | 172Yb | Fluidigm |
| **FoxP3** | 206D & 259D | 173Yb | BioLegend |
| **HLA-DR** | L243 | 174Yb | Fluidigm |
| **CD223 (LAG-3)** | 11C3C65 | 175Lu | Fluidigm |
| **CD127 (IL-7Ra)** | A019D5 | 176Yb | Fluidigm |
| **CD16** | 3G8 | 209Bi | Fluidigm |

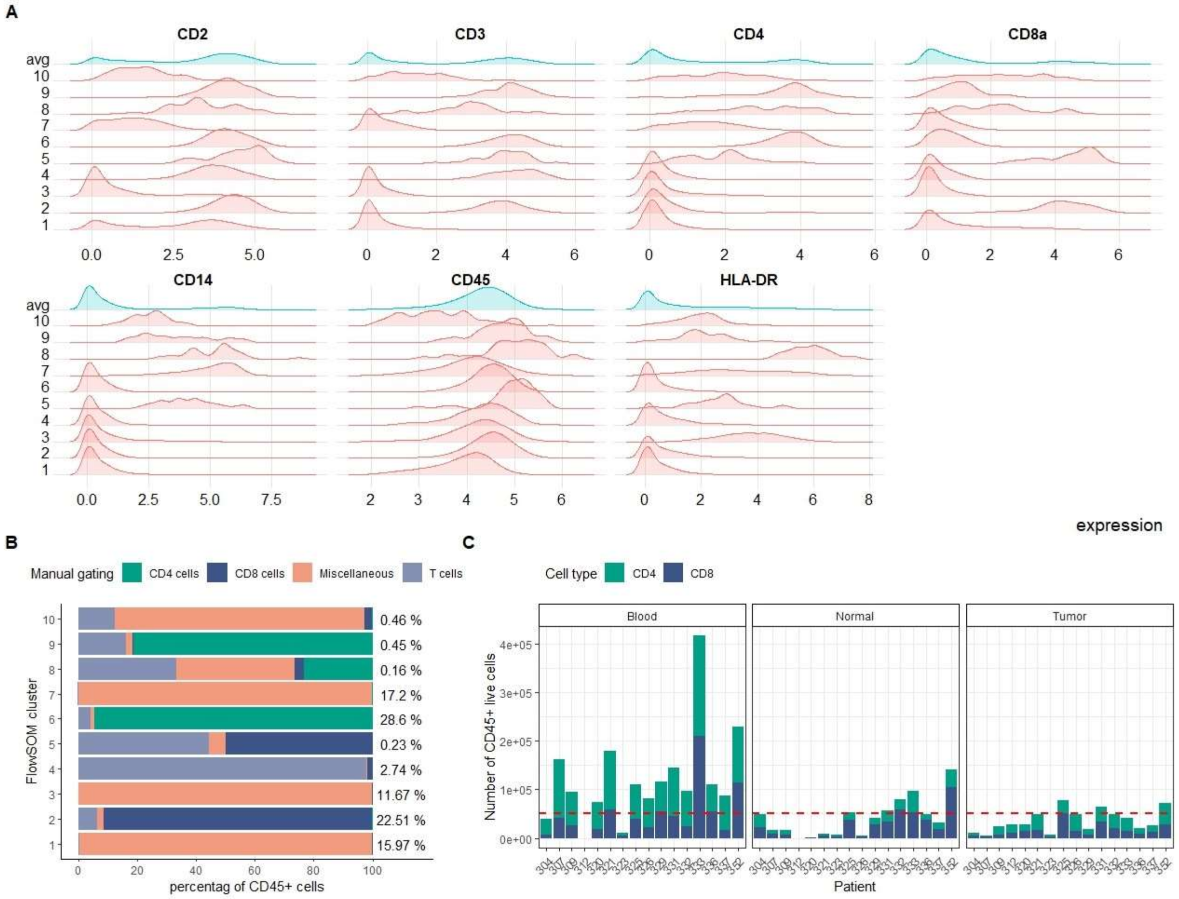


Figure S1. Gating and initial FlowSOM clustering

**(A)** initial clustering for identification of T cells. FlowSOM clustering based on shown lineage markers resulted in 10 clusters. Shown are density plots of normalized marker expression across the clusters **(B)** Percentage of cells shared between manual gating and the 10 clusters. Cluster 2 was identified as CD8 cells and cluster 6 was identified as CD4 cells. On the right, the percentage of CD45+ live cells in the given cluster is indicated **(C)** Total number of CD4 and Cd8 cells in patients' samples across tissues. Downsampling was performed on 50,000 cells or less (indicated with a red line).

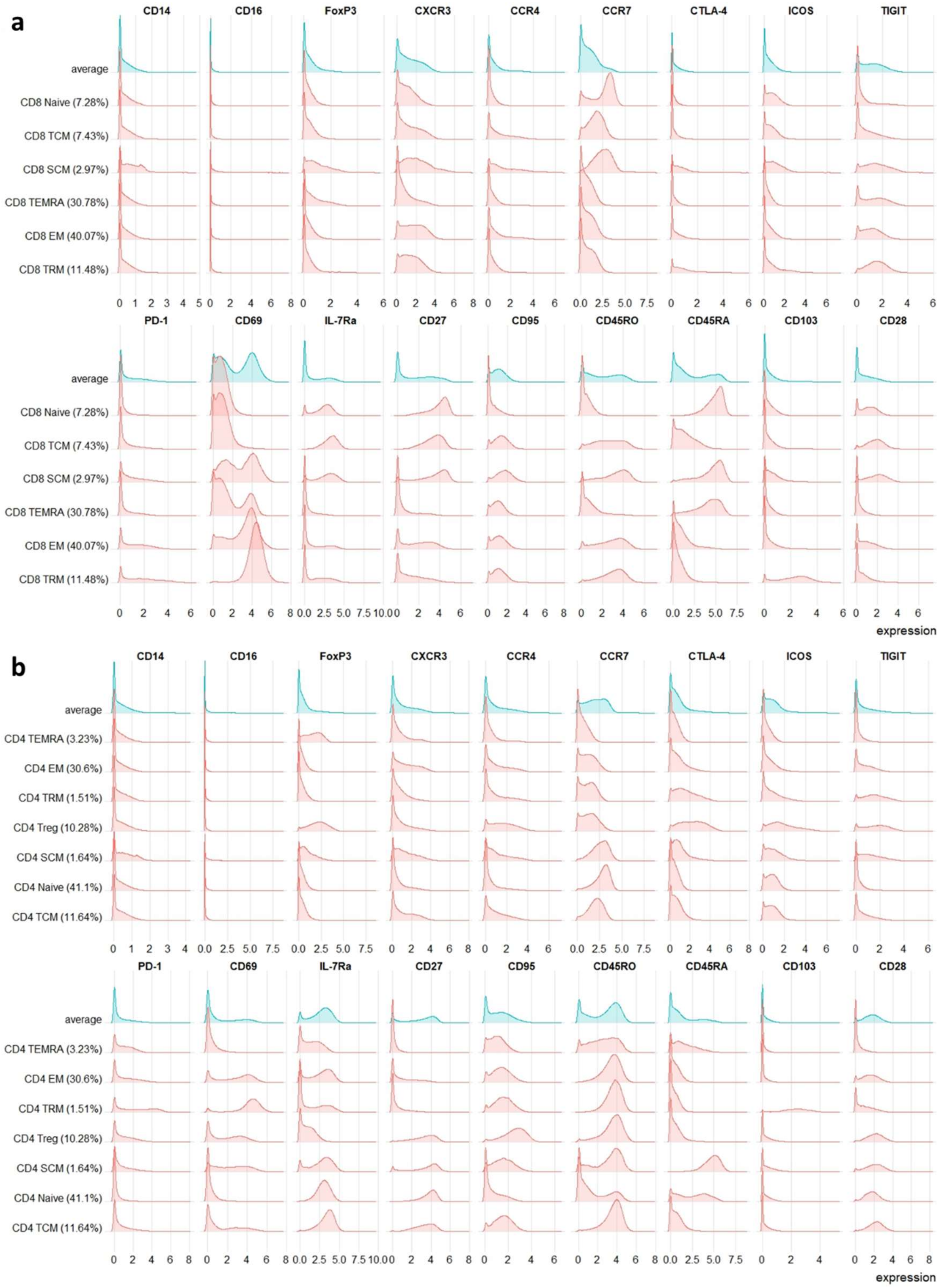


Figure S2. Marker expression in annotated cell types

Density plot of normalized marker expression. Cells were clustered and clusters were annotated based on the marker expression for **(a)** the CD8 dataset and **(b)** the CD4 dataset. The upper blue column (avg) shows the average expression of all CD4+/ CD8+ T cells.

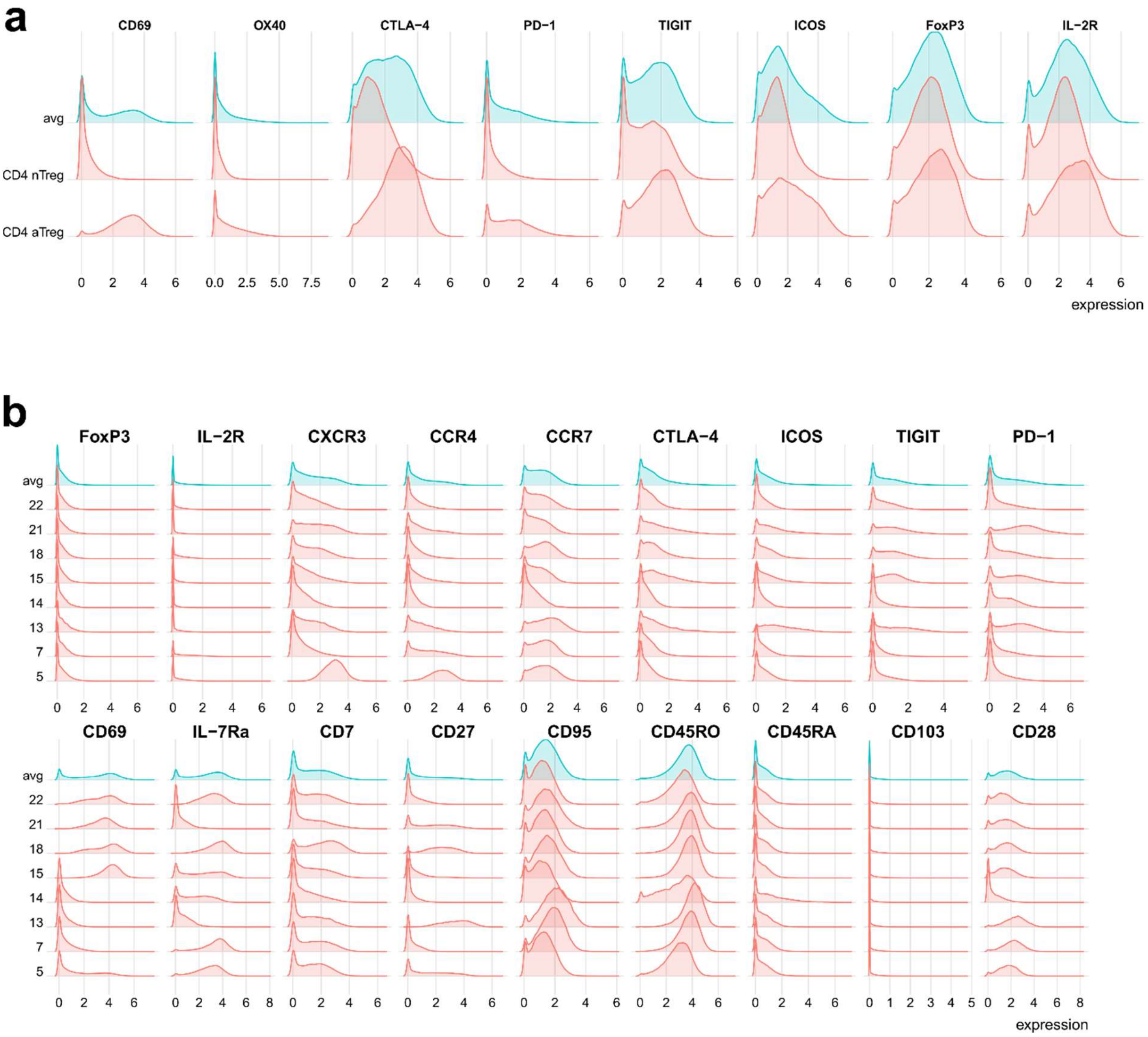


Figure S3. Marker expression of activated Tregs

**(a)** We identified two different clusters of regulatory CD4+ cells and based on their mean marker expression we annotated them into aTregs=activated Tregs and nTregs=none-activated Tregs. Shown are density plots of normalized marker expression. The upper column at the top shows the average expression across all identified Tregs. **(b)** Marker expression of all identified CD4 EM clusters. We observed an increased abundance of cluster 7 in the periphery for patients without lymph node metastasis.

Table S3. Clinical data for patients included in snRNA sequencing

| | Poor | | Good | |
|---|---|---|---|---|
| **Included patients** | 4 | | 3 | |
| **Age in years** | 55.3 | | 69.8 | |
| **RFS in months** | 8 | | 17.7 | |
| **Survival time in months** | 12.7 | | 17.7 | |
| **Grade** | | | | |
| G1 | 0 | 0% | 1 | 33% |
| G2 | 0 | 0% | 1 | 33% |
| G3 | 3 | 75% | 1 | 33% |
| G4 | 0 | 0% | 0 | 0% |
| not known | 1 | 25% | 0 | 0% |
| **Perineural invasion** | | | | |
| positive | 1 | 25% | 0 | 0% |
| negative | 2 | 50% | 3 | 100% |
| not known | 1 | 25% | 0 | 0% |
| **Lymph node invasion** | | | | |
| positive | 4 | 100% | 0 | 0 |
| negative | 0 | 0% | 2 | 100% |
| not known | 0 | 0% | 0 | 0% |
| **Lymphatic or vascular invasion** | | | | |
| positive | 0 | 0% | 1 | 33% |
| negative | 3 | 75% | 2 | 67% |
| not known | 1 | 25% | 0 | 0% |

Table S4. molecular subtypes of tumors determined by next-generation sequencing (NGS)

| Entity | Included in: | Detected variants per patient in gene: |
|---|---|---|
| **iCCA** | CyTOF cohort | **Deletion:**<br>BAP1<br>**Single nucleotide variant:**<br>IDH1 (394C>T)<br>**Unknown significance:**<br>NOTCH1 |
| **iCCA** | CyTOF cohort | **Translocation:**<br>FGFR2 |
| **pCCA** | snRNA cohort (poor) | **Activating:**<br>KRAS (35G>T)<br>**Deletion:**<br>SMAD4<br>**Unknown significance:**<br>ERBB2, ATM |
| **pCCA** | CyTOF cohort<br>snRNA cohort (poor) | **Deletion**:<br>RB, SMARCA4<br>**Unknown significance:**<br>CDKN2A, CDKN2B, ATR, CCNE1 |
| **pCCA** | CyTOF cohort<br>snRNA cohort (good) | **Single nucleotide variant:**<br>KRAS (35 G>A)<br>**Unknown significance:**<br>IDH1, EPHB1, FANCA, KDM5C, MYC, ZNF703 |

Next-generation sequencing data was available for 3 of the pCCA and 2 of the iCCA patients included in the CyTOF and snRNA seq cohorts.

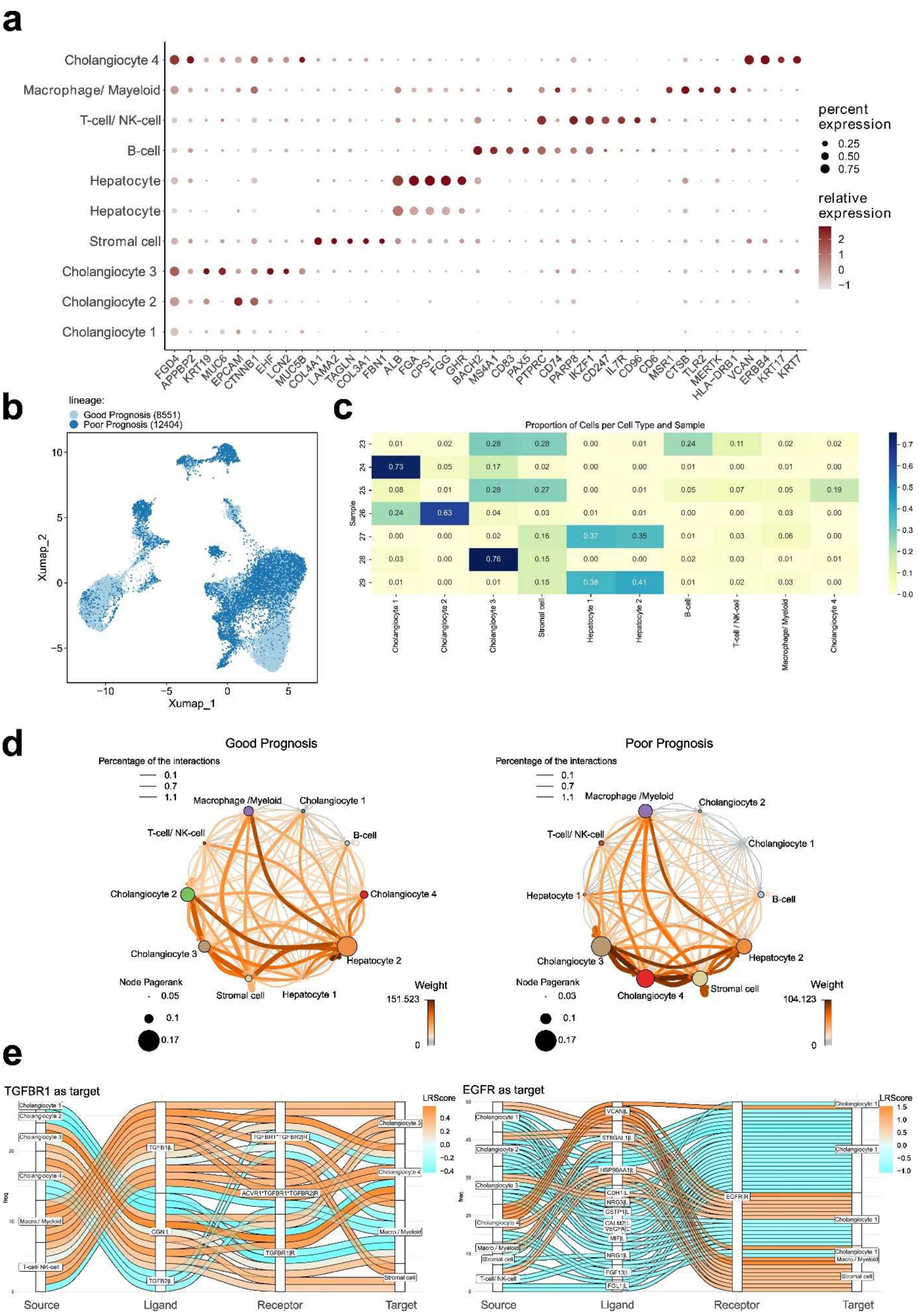
a
Cholangiocyte 4
Macrophage/ Mayeloid
T-cell/ NK-cell
B-cell
Hepatocyte
Hepatocyte
Stromal cell
Cholangiocyte 3
Cholangiocyte 2
Cholangiocyte 1
FGD4 APPBP2 KRT19 MUC6 EPCAM CTNNB1 EHF LCN2 MUC5B COL4A1 LAMA2 TAGLN COL3A1 FBN1 ALB FGA CPS1 FGG GHR BACH2 MS4A1 CD83 PAX5 PTPRC CD74 PARP8 IKZF1 CD247 IL7R CD96 CD6 MSR1 CTSB TLR2 MERTK HLA-DRB1 VCAN ERBB4 KRT17 KRT7
percent expression
0.25
0.50
0.75
relative expression
2
1
0
−1
b
lineage:
Good Prognosis (8551)
Poor Prognosis (12404)
Xumap_2
Xumap_1
c
Proportion of Cells per Cell Type and Sample
Sample
23: 0.01 0.02 0.28 0.28 0.00 0.01 0.24 0.11 0.02 0.02
24: 0.73 0.05 0.17 0.02 0.00 0.00 0.01 0.00 0.01 0.00
25: 0.08 0.01 0.28 0.27 0.00 0.01 0.05 0.07 0.05 0.19
26: 0.24 0.63 0.04 0.03 0.01 0.01 0.00 0.00 0.03 0.00
27: 0.00 0.00 0.02 0.16 0.37 0.35 0.01 0.03 0.06 0.00
28: 0.03 0.00 0.76 0.15 0.00 0.00 0.02 0.03 0.01 0.01
29: 0.01 0.00 0.01 0.15 0.38 0.41 0.01 0.02 0.03 0.00
Cholangiocyte 1 Cholangiocyte 2 Cholangiocyte 3 Stromal cell Hepatocyte 1 Hepatocyte 2 B-cell T-cell / NK-cell Macrophage/ Myeloid Cholangiocyte 4
d
Good Prognosis
Poor Prognosis
Percentage of the interactions
0.1
0.7
1.1
Macrophage /Myeloid
Cholangiocyte 1
Cholangiocyte 2
T-cell/ NK-cell
B-cell
Cholangiocyte 4
Cholangiocyte 3
Hepatocyte 1
Hepatocyte 2
Stromal cell
Node Pagerank
0.05
0.03
0.1
0.17
Weight
151.523
104.123
0
e
TGFBR1 as target
EGFR as target
LRScore
Source
Ligand
Receptor
Target
TGFB1|L
CGN|L
TGFB2|L
TGFBR1*TGFBR2|R
ACVR1*TGFBR1*TGFBR2|R
TGFBR1|R
VCAN|L
ST6GAL1|L
HSP90AA1|L
CDH1|L
NRG3|L
GSTP1|L
CALM2|L
VEGFA|L
MIF|L
NRG1|L
FGF13|L
FGL1|L
EGFR|R
Macro./ Myeloid
Stromal cell
T-cell/ NK-cell

Figure S4. snRNA clusters and cell communication

Data from snRNA sequencing was clustered into 9 major cell clusters and annotated based on marker expression. **(a)** Expression of marker genes used for cell type annotation. **(b)** UMAP dimensionality reduction of single-cell data, colored by prognosis group. **(c)** Proportion of each cell type by sample. **(d)** Cell-cell communication network for patients with good prognosis (left) and poor prognosis (right). **(e)** Sankey plots visualize the top interactions of the target genes EGFR and TGFBR1 predicted by the cell-cell communication analysis. Shown are Ligand receptor pairs as well as the cell types expressing the ligand (source) and receptor (target).

Table S5. Top Ligand receptor pairs for CrosstalkeR analysis

| Ligand cluster | Receptor cluster | Ligand | Receptor | LRScore | lr_type |
|---|---|---|---|---|---|
| Cholangiocyte4 | SCHSC | VCAN | EGFR | 1.530624734 | up |
| Cholangiocyte4 | SCHSC | VCAN | ITGB1 | 1.411956534 | up |
| Cholangiocyte4 | SCHSC | C3 | NRP1 | 1.353639209 | up |
| Cholangiocyte4 | SCHSC | VCAN | CD44 | 1.314084793 | up |
| Cholangiocyte4 | SCHSC | C3 | CD46 | 1.2500376 | up |
| Cholangiocyte4 | SCHSC | C3 | LRP1 | 1.23433813 | up |
| Cholangiocyte4 | SCHSC | COMP | ITGA5 | 1.183703367 | up |
| Cholangiocyte4 | SCHSC | SHANK2 | CFTR | 1.148588111 | up |
| Cholangiocyte4 | SCHSC | COMP | CD47 | 1.071311731 | up |
| Cholangiocyte4 | SCHSC | NAMPT | INSR | 1.00965035 | up |
| Cholangiocyte4 | Macro-Myeloid | C3 | ITGAX | 1.47803287 | up |
| Cholangiocyte4 | Macro-Myeloid | VCAN | TLR2 | 1.532430001 | up |
| Cholangiocyte4 | Macro-Myeloid | C3 | ITGAX | 1.47803287 | up |
| Cholangiocyte4 | Macro-Myeloid | VCAN | CD44 | 1.43987253 | up |
| Cholangiocyte4 | Macro-Myeloid | VCAN | ITGB1 | 1.318776884 | up |
| Cholangiocyte4 | Macro-Myeloid | C3 | NRP1 | 1.299391232 | up |
| Cholangiocyte4 | Macro-Myeloid | VCAN | ITGA4 | 1.277297876 | up |
| Cholangiocyte4 | Macro-Myeloid | VCAN | EGFR | 1.274140735 | up |
| Cholangiocyte4 | Macro-Myeloid | C3 | LRP1 | 1.263180149 | up |
| Cholangiocyte4 | Macro-Myeloid | C3 | CD46 | 1.25411593 | up |
| Cholangiocyte4 | Macro-Myeloid | SHANK2 | CFTR | 1.171654025 | up |
| Macro-Myeloid | Cholangiocyte4 | TGFB1 | LPP | 1.353521378 | up |
| Macro-Myeloid | Cholangiocyte4 | C3 | NRP1 | 0.869658725 | up |
| Macro-Myeloid | Cholangiocyte4 | MAML2 | NOTCH3 | 0.866072468 | up |
| Macro-Myeloid | Cholangiocyte4 | ADAM17 | ERBB4 | 0.859836099 | up |
| Macro-Myeloid | Cholangiocyte4 | ALCAM | NRP1 | 0.759517062 | up |
| Macro-Myeloid | Cholangiocyte4 | THBS1 | PTPRJ | 0.706997703 | up |
| Macro-Myeloid | Cholangiocyte4 | ADAM10 | MET | 0.682669852 | up |
| Macro-Myeloid | Cholangiocyte4 | ADAM17 | MET | 0.679590567 | up |
| Macro-Myeloid | Cholangiocyte4 | SEMA4D | MET | 0.607548235 | up |
| Macro-Myeloid | Cholangiocyte4 | ADAM10 | IL6R | 0.58661754 | up |

The top 10 Ligand receptor pairs for the interaction between Cholangiocyte cluster 4 with Stromal/ Hepatic stellate cells (SCHSC) and between cholangiocyte 4 and Macrophages and myeloid cells (macro-myeloid) in both directions. Filtering was done based on log Fold change of the Ligand receptor pairs.

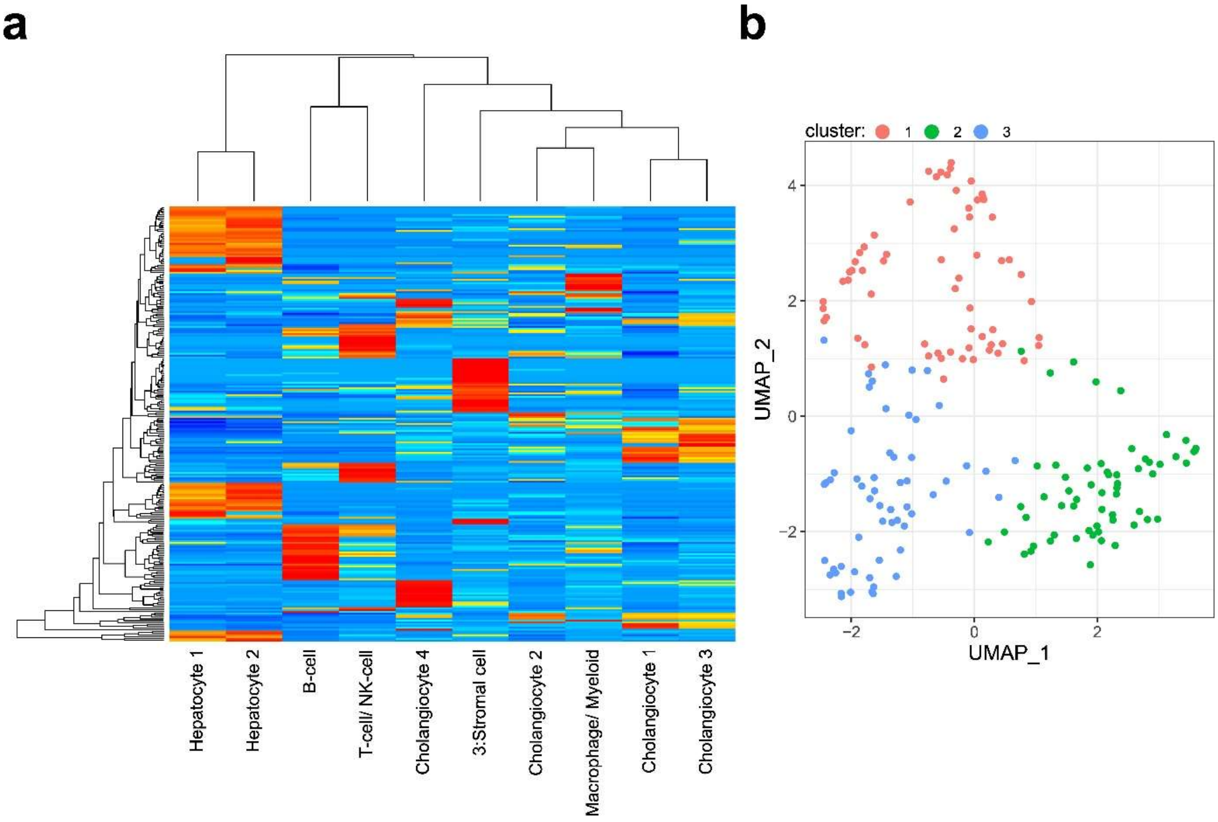


Figure S5. Signature matrix is used for cellular deconvolution and clustering based on the deconvolution results

**(a)** We used cibersortx to construct a signature matrix based on our 10 clusters identified in the snRNA data. The signature matrix was used to predict cellular fractions in public transcriptomic data from 182 eCCA patients. **(b)** Relative fractions within the samples were used as input for dimensionality reduction and clustering resulting in 3 distinct clusters.
